# APOE4 Lowers Energy Expenditure and Impairs Glucose Oxidation by Increasing Flux through Aerobic Glycolysis

**DOI:** 10.1101/2020.10.19.345991

**Authors:** Brandon C. Farmer, Holden C. Williams, Nicholas Devanney, Margaret A. Piron, Grant K. Nation, David J. Carter, Adeline E. Walsh, Rebika Khanal, Lyndsay E. A. Young, Jude C. Kluemper, Gabriela Hernandez, Elizabeth J. Allenger, Rachel Mooney, J. Anthony Brandon, Vedant A. Gupta, Philip A. Kern, Matthew S. Gentry, Josh M. Morganti, Ramon C. Sun, Lance A. Johnson

## Abstract

Cerebral glucose hypometabolism is consistently observed in individuals with Alzheimer’s disease (AD), as well as in young cognitively normal carriers of the E4 allele of Apolipoprotein E (APOE), the strongest genetic predictor of late-onset AD. While this clinical feature has been described for over two decades, the mechanism underlying these changes in cerebral glucose metabolism remains a critical knowledge gap in the field. Here, we undertook a multi-omic approach by combining single-cell RNA sequencing (scRNAseq) and stable isotope resolved metabolomics (SIRM) to define a metabolic rewiring across astrocytes, brain tissue, mice, and human subjects expressing APOE4. Single-cell analysis of brain tissue from mice expressing human APOE revealed E4-associated decreases in genes related to oxidative phosphorylation, particularly in astrocytes. This shift was confirmed on a metabolic level with isotopic tracing of ^13^C-glucose in E4 mice and astrocytes, which showed decreased pyruvate entry into the TCA cycle and increases in lactate synthesis. Metabolic phenotyping of E4 astrocytes showed elevated glycolytic activity, decreased oxygen consumption, blunted oxidative flexibility, and a lower rate of glucose oxidation in the presence of lactate. Together, these cellular findings suggested an E4 associated increase in aerobic glycolysis (i.e. the Warburg effect). To test whether this phenomenon translated to APOE4 humans, we analyzed the plasma metabolome of young and middle-aged human participants with and without the E4 allele, and used indirect calorimetry to measure whole body oxygen consumption and energy expenditure. In line with data from E4-expressing mice, young female E4 carriers showed a striking decrease in energy expenditure compared to non-carriers. This decrease in energy expenditure was primarily driven by a lower rate of oxygen consumption, and was exaggerated following a dietary glucose challenge. Further, the stunted oxygen consumption was accompanied by markedly increased lactate in the plasma of E4 carriers, and a pathway analysis of the plasma metabolome suggested an increase in aerobic glycolysis. Together, these results suggest astrocyte, brain and system-level metabolic reprogramming in the presence of APOE4, a ‘Warburg like’ endophenotype that is observable in young humans decades prior to clinically manifest AD.

## Introduction

The E4 allele of Apolipoprotein E (*APOE*) confers more risk (up to 15 fold) for the development of late-onset Alzheimer’s disease (AD) than any other gene ^1,2^. While E4 is a strong contributor to late-onset AD risk, the effect is even greater in females ^3^. Female E4 carriers have an increased odds ratio for AD ^4^, increased incidence of AD ^5^, elevated hazard ratio for conversion to mild cognitive impairment ^6^, increased CSF tau ^7^, and reduced hippocampal volume ^8^, compared to male E4 carriers. To date, studies investigating the mechanism by which E4 and sex increase disease risk have primarily focused on the important associations of E4 with the neuropathological hallmarks of AD – i.e. the increased amyloid load seen in E4 carriers ^9,10^ and the *APOE*-dependence of tau propagation ^11,12^.

Alternatively, investigating E4 carriers who have not yet developed neuropathology may provide insight into early E4 mechanisms and unveil additional therapeutic targets for the prevention of AD. For example, an early and consistent biological hallmark of AD is cerebral glucose hypometabolism as observed by ^18^F-fluorodeoxyglucose positron emission tomography (FDG-PET) imaging ^13–15^. Interestingly, E4 carriers also display an “AD-like” pattern of decreased glucose metabolism by FDG-PET long before clinical symptomology ^16,17^. Since glucose hypometabolism occurs early in AD and early in E4 carriers, it may represent a critical initial phase of AD pathogenesis that predisposes individuals to subsequent symptomology.

Beyond this FDG-PET finding, it is not clear if *APOE* has other discernable metabolic effects in pre-cognitively impaired young people, and clinical research focused on how *APOE* may regulate metabolism outside of the brain is limited ^18^. Most studies have utilized a targeted replacement mouse model of *APOE* in which the murine *Apoe* alleles are replaced by the human orthologs ^19,20^. For example, several studies have found E4 mice to exhibit increased susceptibility to insulin resistance, and one report characterized E4 mice as deficient in extracting energy from dietary sources ^21–23^. While these preclinical studies have been critical to our understanding of E4-associated impairments in glucose metabolism, the mechanism underlying these changes, and the extent to which systemic glucose metabolism is regulated by *APOE* in young healthy humans, remain largely unknown.

In the current study, we combined single-cell RNA sequencing (scRNAseq) and stable isotope resolved metabolomics (SIRM) to define a metabolic shift toward aerobic glycolysis across astrocytes, brain tissue, mice, and human subjects expressing APOE4. We highlight an astrocyte-directed shift in gene expression away from oxidative phosphorylation in the brains of mice expressing human E4, and confirm this metabolic reprogramming through the use of isotopic tracing of ^13^C-glucose in both E4 mice and astrocytes. To test whether this phenomenon translated to APOE4 humans, we used indirect calorimetry to measure whole body oxygen consumption and energy expenditure in young and middle-aged human participants with and without the E4 allele. Strikingly, young female E4 carriers showed a significant decrease in resting energy expenditure compared to non-carriers, a decrease driven primarily by reductions in oxygen consumption. Interestingly, this stunted oxygen consumption was exaggerated following a dietary glucose challenge and was accompanied by markedly increased lactate in the plasma of E4 carriers. Together, these results suggest astrocyte, brain and system-level metabolic reprogramming in the presence of APOE4, a pro-glycolytic shift that is observable in young women decades prior to clinically manifest AD.

## Materials and Methods

### Clinical research study design

The study objectives were to i) determine if *APOE* genotype influences peripheral and cerebral metabolism in young cognitively normal human subjects, and if so, ii) elucidate potential mechanisms using mouse and cell models of human *APOE*. For the clinical research study, healthy volunteers between 18-65 were prescreened for diagnoses that may affect cognitive function (ex. stroke, Parkinson’s), metabolic diseases (diabetes), alcoholism, drug abuse, chronic major psychiatric disorders, medications that interfere with cognition (narcotic analgesics, anti-depressants), medications that interfere with EE (stimulants, beta-blockers) and vision or hearing deficits that may interfere with testing. The prescreening checklist with a full list of medications and conditions excluded for can be found in the supplemental materials (Extended Data Table 5). Eligible candidates were brought in for informed consent after a 12-hour fast in which subjects were asked not to exercise and to abstain from everything except for water. We employed a power analysis based on a feasibility study, and the required sample size per group for a power level of 0.9 was calculated to be n = 30 per “group” (i.e. E2+, E3/E3 and E4+), for a total of 90 subjects. To account for potential biological outliers, non-consenting subjects, and post-recruitment exclusion criteria being met, we recruited a total of 100 individuals for this observational study. Prior to unblinding to *APOE* genotype, individuals who had IC values more than 2 standard deviations from the mean were excluded from analysis, leaving 94 individuals for analysis. As we were primarily interested in *APOE* effects in young individuals, we stratified our sample population into a young cohort (under 40 years old) and a middle-aged cohort (40-65 years old). We chose 40 as the age-cutoff based on a meta-analysis of *APOE* genotype and AD-risk which found the E4 effect on disease to be observable in individuals 40 and over ^4^. Data acquisition was blinded as *APOE* genotypes were determined after the study. Body mass index (BMI), waist to hip ratio, and blood pressure were first recorded. Thereafter, participants were fitted with an airtight mask that was connected to an MGC Diagnostics Ultima CPX metabolic cart which measures VO_2_, VCO_2_, and respiratory rate. EE is defined as the amount of energy an individual uses to maintain homeostasis in kcal per day, and can be calculated using the Weir equation (EE = 1.44 (3.94 VO_2_ + 1.11 VCO_2_) ^24^. EE is composed of the resting energy expenditure (REE), the thermic effect of feeding (TEF), and activity related energy expenditure (AEE). In motionless and fasted humans, EE is equivalent to the REE since the TEF and AEE have been controlled for. Participants were instructed to remain motionless and to refrain from sleep for 30 minutes as data was gathered. All testing occurred between 8:30-11:30 am in a temperature controlled (20–22 °C) out-patient research unit (Center for Clinical and Translational Science, University of Kentucky). Body temperature was taken periodically via temporal thermometer to ensure thermostasis and provide intermittent stimulation to ensure wakefulness. After the resting period came a 30 minute cognitive test period. We then introduced a novel-image-novel-location (NINL) object recognition test consisting of a series of images which participants were later asked to recall. This test has been shown previously to study *APOE* allele effects on cognition ^25^. After the cognitive test period, a blood draw was taken via venipuncture and placed on ice. Participants then consumed a sugary milk drink consisting of 50g of sugar dissolved in whole milk. The drink was consumed within a two minute time span. The mask was then refitted and participants were instructed to again remain motionless for 30 minutes for data collection. Data from the first 5 minutes of the study time periods were excluded to allow a five minute steady state adjustment ^26,27^. After the glucose challenge, participants provided a second blood sample (~45 minutes after the initial blood draw). Participants then exited the study and were compensated for their participation. This study was approved by the University of Kentucky Institutional Review Board (#48365) and was listed as Clinical Trial #NCT03109661.

### *APOE* Genotyping

*APOE* genotype was determined by extracting genomic DNA from participants’ blood samples using a GenElute Blood Genomic DNA Kit (Sigma). After confirming concentration and quality by Nanodrop, *APOE* genotype was determined using PCR with TaqMan assay primers for the two allele-determining SNPs of *APOE*: rs7412 and rs429358 (Thermo). Positive controls for the six possible *APOE* genotypes were included with each assay.

### Plasma metabolomics and GCMS Sample Preparation

Plasma was separated from blood by centrifugation at 2500 x *g* for 10 minutes at 4°C, and stored in 200uL aliquots at −80°C until further use. Upon thawing, ice cold 100% methanol solution containing 40nM L-norvaline (internal standard) was added to 80 μl of plasma and kept on ice for 20 minutes with regular vortexing. The solution was then centrifuged for 10 minutes (14,000 rpm, 4°C). Supernatant containing polar metabolites was removed to a new tube and kept at −80°C until prepped for GCMS analysis. Polar metabolites were thawed on ice then dried under vacuum. The dried pellet was dissolved in 50 μL methoxyamine HCl-pyridine (20 mg/ml) solution and heated 60 minutes at 60°C. Following heating, samples were transferred to v-shaped glass chromatography vials and 80 μl of MTSFA + 1% TMCS (Thermo Scientific) was added. Samples were then heated for 60 minutes at 60°C, allowed to cool to room temperature, and then analyzed via GCMS with parameters as previously described ^28^. Briefly, a GC temperature gradient of 130°C was held for 4 minutes, rising at 6°C/min to 243°C, rising at 60°C/min to 280°C and held for two minutes. Electron ionization energy was set to 70 eV. Scan and full scan mode used for metabolite analysis, spectra were translated to relative abundance using the Automated Mass Spectral Deconvolution and Identification System (AMDIS) software with retention time and fragmentation pattern matched to FiehnLib library with a confidence score of >80. Chromatograms were quantified using Data Extraction for Stable Isotope-labelled metabolites (DExSI) with a primary ion and two or more matching qualifying ions. Metabolite quantification was normalized to relative abundance of internal standard (L-norvaline), brain and cell data also normalized to protein concentration. Metabolomics data was analyzed using the web-based data processing tool Metaboanalyst ^29^. Metabolites significantly altered by *APOE* genotype and/or time point were defined by ANOVA and subsequent false discovery rate cutoff of < 0.05. All identified metabolites for which >75% of participants had a measurable concentration were included, and missing values were estimated with an optimized random forest method ^30^. For the pathway impact analysis, the parameters were set to ‘global test’ and ‘Relative-betweenness Centrality’, a node centrality measure which reflects metabolic pathway ‘hub’ importance. For enrichment analyses, parameters were set to “Pathway-associated metabolite sets (SMPDB)”, a library that contains 99 metabolite sets based on normal human metabolism. For both pathway and impact analyses, only metabolic pathways with 3+ metabolites represented in our data set were included, and a false discovery rate cutoff of <0.05 was utilized.

### Mice and metabolic phenotyping

Mice expressing human *APOE* display many of the phenotypic characteristics observed in humans including several metabolic variations noted in epidemiological studies ^31–33^. In this “knock-in” model, the mouse *Apoe* locus is targeted and replaced with the various human *APOE* alleles, thereby remaining under control of the endogenous mouse *Apoe* promoter and resulting in a physiologically relevant pattern and level of human *APOE* expression ^17,34–38^. Mice used in this study were homozygous for either the human E3 or E4 alleles, aged 2-4 months (young) and group housed in sterile micro-isolator cages (Lab Products, Maywood, NJ), and fed autoclaved food and acidified water *ad libitum*. Animal protocols were reviewed and approved by the University of Kentucky Institutional Animal Use and Care Committee. Human E3 and E4 mice were evaluated by indirect calorimetry (TSE Systems, Chesterfield, MO). Mouse body composition was measured using EchoMRI (Echo Medical Systems, Houston, TX) the morning prior to being singly housed in the indirect calorimetry system. Mice were acclimated to singly housed cage conditions for one week prior to beginning data recording. After five days on standard chow diet (Teklad Global 18% protein rodent diet; 2018; Teklad, Madison, WI), mice were fasted overnight before being introduced to a high carb diet (Open Source Diets, Control Diet for Ketogenic Diet with Mostly Cocoa Butter, D10070802) for five days. Mice were monitored for O_2_ consumption, CO_2_ production, movement, and food and water consumption. Chambers were sampled in succession and were reported as the average of 30 minute intervals in reference to an unoccupied chamber. To negate the effects of activity on EE readouts, we chose to only analyze the light cycles of the mice where activity, and feeding, is minimal. The EE then becomes analogous to a “resting” EE similar to the resting period in the human study and differences observed are likely due to basal metabolic rate differences instead of confounding factors such as feeding and activity ^39^.

### Cell culture

Primary astrocytes were isolated from postnatal day 0-4 pups of mice homozygous for E3 or E4. The brain was surgically excised and meninges were removed from cortical tissue in cold DMEM. Tissue from pups of the same genotype was pooled and coarsely chopped to encourage suspension. Tissue homogenates were incubated in serum free DMEM with 0.25% trypsin and DNAse for 30 min with gentle shaking. Cell suspension was then filtered through 40 μm strainer and spun for 5 min at 1100 x *g*. Suspended primary cells were then plated in a poly-lysine coated plate and allowed to grow to confluence in Advanced DMEM (Gibco) with 10% FBS. Immortalized astrocytes were derived from targeted replacement mice expressing human *APOE* alleles (kind gift from Dr. David Holtzman). These immortalized cell lines secrete human ApoE in HDL-like particles at equivalent levels to primary astrocytes from targeted replacement *APOE* knock-in mice and have been relied upon for studies of *APOE*’s role in astrocyte metabolism by several groups ^40–42^. Cells were maintained in Advanced DMEM (Gibco) supplemented with 1mM sodium pyruvate, 1X Geneticin, and 10% fetal bovine serum unless otherwise noted.

### Single-cell RNA sequencing

Brain tissues were processed for creating single cell suspensions as previously described (Early et al., 2020). Briefly, E3 and E4 mice (pooled n=3 per genotype) were anesthetized via 5.0% isoflurane before exsanguination and transcardial perfusion with ice-cold Dulbecco’s phosphate buffered saline (DPBS; Gibco # 14040133). Following perfusion, brains were quickly removed and whole hemispheres sans brainstem and cerebellum were quickly minced using forceps on top of an ice-chilled petri dish. Minced tissue from the 3 pooled hemispheres per genotype were immediately transferred into gentleMACS C-tube (Miltenyi #130-093-237) containing Adult Brain Dissociation Kit (ADBK) enzymatic digest reagents (Miltenyi #130-107-677) prepared according to manufacturer’s protocol. Tissues were dissociated using the “37C_ABDK” protocol on the gentleMACS Octo Dissociator instrument (Miltenyi #130-095-937) with heaters attached. After tissue digestion, cell suspensions were processed for debris removal and filtered through 70μm mesh cell filters following the manufacturer’s suggested ABDK protocol. The resultant suspension was filtered sequentially two more times using fresh 30μm mesh filters. Cell viability was checked using AO/PI viability kit (Logos Biosystems # LGBD10012) both cell suspensions were determined to have >88% viable cells. Following viability and counting, cells were diluted to achieve a concentration of ~1000 cells/100uL. The diluted cell suspensions were loaded onto the 10x Genomics Chromium Controller. Each sample was loaded into a separate channel on the Single Cell 3’ Chip and libraries were prepared using the Chromium v3 Single Cell 3’ Library and Gel Bead Kit (10x Genomics). Final library quantification and quality check was performed using BioAnalyzer (Agilent), and sequencing performed on a NovaSeq 6000 S4 flow cell, 150bp Paired-End sequencing (Novogene). Raw sequencing data was de-multiplexed and aligned using Cell Ranger (10x Genomics), and further processed using Partek software. To remove likely multiplet and dead cells, cells were discarded if they had total read counts less than 50 or greater than 50000 UMIs, or mitochondrial read counts more than 30%. UMAP projections were visualized with 20 principal components. Clusters were assigned to cell types using known marker genes. Two small clusters (<250 cells) were removed from downstream analysis due to suspected doublets/triplets based on positive gene expression of multiple cell-specific gene markers (astrocytes, microglia, mural cells and/or endothelial cells). The final dataset consisted of a total of 18,167 cells (8,216 and 9,951 cells from E3 and E4, respectively) that passed quality control thresholds.

### Glucose tracing *in vivo*

Female TR mice homozygous for E3 or E4 (12-13 month) were fasted for 2-3 hours then, via oral gavage, administered 250 μL [U-^13^C] glucose solution at a concentration of 2 g/kg of body weight based on average cohort bodyweight. 45 minutes following gavage, mice were euthanized by cervical dislocation, brains were removed and quickly washed twice in PBS, once in H_2_O then frozen in liquid N_2_. Tissues were kept at −80°C until ground under liquid N_2_ using a Freezer/Mill Cryogenic Grinder (SPEX SamplePrep model 9875D). Approximately 60 mg of tissue was placed in a 1.5 mL tube then 1 mL extraction buffer (50% methanol, 20nM norvaline) was added followed by a brief vortex and placed on ice for 20 min, briefly vortexed every 5 min. Samples were then centrifuged at 14,000 rpm, 4°C for 10 min. The supernatant containing polar metabolites was removed to a new tube and kept at −80°C until prepped for GCMS. The resulting pellet was re-suspended in RIPA buffer (Sigma) and protein concentration was measured with BCA kit (Pierce) for normalization.

### Glucose metabolism assays

For glucose oxidation assays, astrocytes were plated in a 24-well plate at 300,000 cells/well with 500 μL of maintenance media (Advanced DMEM, 10% FBS, 1% sodium pyruvate, 0.4% Geneticin) and incubated at 5% CO_2_ and 37°C and allowed to grow to confluence for 24 hours. Using a previously published protocol ^43^, cells were then incubated with 1 μCi/mL [U-^14^C] glucose in maintenance media (25mM glucose) or starvation media (same as maintenance except 0mM glucose) for 3 hours. Buffered ^14^CO_2_ in the media was then liberated by addition of 1 M perchloric acid and captured on a filter paper disc pre-soaked with 1N sodium hydroxide using airtight acidification vials. Radioactivity of the filter paper was measured in a Microbeta 2 Scintillation Counter (Perkin Elmer) after addition of 3 mL Ultima-Gold Scintillation Fluid. For glucose tracing in primary astrocytes, cells were plated in a 6-well plate at 600,000 cell/well in astrocyte growth media (Advanced DMEM, 10% FBS, 1% sodium pyruvate, 1% penicillin-streptomycin) and incubated at 5% CO_2_ and 37°C. After 48 hours, growth media was replaced with tracer media (Glucose-free DMEM containing 10% dialyzed FBS, 10mM [U-^13^C] glucose) and incubated under previous conditions for 24 hours at which time quenching and metabolite extraction were carried out as follows: Plates were retrieved from incubator and placed on ice, tracer media removed and wells washed once with ice-cold PBS. Immediately following washing, 1 mL of ice-cold extraction buffer (50% methanol, 20nM norvaline) was added to quench enzymatic activity and plates were placed at −20°C for 10 min. Cellular contents were then scraped with a cell-scraper in extraction buffer and collected into 1.5 mL and tubes placed on ice for 20 min with regular vortexing. Samples were then centrifuged at 14,000 rpm, 10 min, 4°C after which supernatant containing polar metabolites were removed to a new tube and frozen at −80°C until prepped for GCMS analysis. The resulting pellet was re-suspended in RIPA buffer (Sigma) and protein concentration was measured with BCA kit (Pierce) for normalization.

### Mitochondrial respiration assays

Astrocytes were plated at 40,000 cells/well in maintenance media and grown to confluence for 24 hours. The following day media was replaced with assay running media (Seahorse XF Base Medium, 1mM pyruvate, 2mM glutamine, and 10mM glucose) and after 1 hour oxygen Consumption rate (OCR) and extracellular acidification rate (ECAR) were measured using a Seahorse 96XF instrument as previously described ^44^. Manufacturer protocols were followed for the glycolysis stress test assay and Mito fuel flex assay. Briefly, the glycolysis stress test assesses the ability of cells to respond to challenging conditions by increase the rate of glycolytic activity. Glycolytic capacity refers to the glycolytic response to energetic demand from stress (Glycolytic capacity = ECAR post-oligomycin – Baseline ECAR) while glycolytic reserve refers to the capacity available to utilize glycolysis beyond the basal rate (Glycolytic reserve = ECAR post-oligomycin – ECAR post-glucose). The Mito Fuel Flex assay assesses mitochondrial energy consumption by measuring respiration in the presence or absence of fuel pathway inhibitors. The following equations were used in the calculations of mitochondrial flexibility parameters: Dependency (%) = [(Baseline OCR - Target inhibitor OCR)/ (Baseline OCR - All inhibitors of OCR)] x 100%. Capacity (%) = 1 / [(Baseline OCR - Other two inhibitors of OCR)/(Baseline OCR - All inhibitors of OCR)] x 100%. Flexibility (%) = Capacity (%) - Dependency (%).

### Statistical analysis

All results are reported as mean +/− SEM unless otherwise stated. For comparisons between two groups, an unpaired two-tailed Student’s *t*-test was used. For pair-wise comparison of two time points a paired two-tailed Student’s *t*-test was used. One-way analysis of variance (ANOVA) was used for comparing multiple groups followed by Sidak’s multiple comparisons test. Two-way ANOVA with repeated measures was used for time course analysis. Pearson r correlation test was used for correlative analysis. For dependent variables with categorical independent variables we analyzed covariance (ANCOVA) to assess collinearity. *P*<0.05 was considered significant.

## Results

### Single-cell RNA sequencing highlights a role for APOE4 in astrocyte oxidative phosphorylation and glycolysis

Given the outsized role of *APOE* in modulating AD risk, we first undertook an unbiased survey of E4 effects in various cell types by performing single cell RNA sequencing (scRNA-seq) on brain tissue from mice expressing human E3 or E4. To visualize and identify cell populations with distinct transcriptional signatures, we performed a Uniform Manifold Approximation and Projection (UMAP) on a total of 18,167 cells (E3 8,216; E4 9,951) from pooled (n=3) whole brain tissue (Fig. 1a; Supplemental Fig. 1a). We then used a list of established marker genes to assign cluster identity (Fig. 1b), including four clusters that highly expressed *Aldoc, Aqp4, Gja1* and *Aldh1l1,* which we assigned as astrocytes (Fig. 1b, blue; Supplemental Fig. 1b). Notably, these astrocyte clusters showed both the highest expression of *APOE* (Fig. 1c) and the highest cumulative expression of a list of 39 genes directly involved in glycolysis (Fig. 1d). When we performed a sub-UMAP on only astrocytes, the cells clustered into eight unique subpopulations with distinct transcriptional signatures (Fig. 1e; Supplemental Table 1). Interestingly, *APOE* expression was higher in E4 astrocytes, an effect primarily driven by clusters 1, 2, 3 and 5 (Supplemental Fig. 2). As expected based on previous bulk sequencing studies of human APOE mice and *APOE* genotyped human brain tissue, a number of other differentially expressed genes (DEGs) were noted between E4 vs E3 cells, including 562 DEGs specifically in astrocytes (Fig. 3f). Notably, gene ontology (GO) analyses of all cells underscored a number of metabolic processes, including several mitochondrial related GO terms (Fig. 1g). In particular, pathway enrichment analyses specifically highlighted “Alzheimer’s disease” and “oxidative phosphorylation” as top hits in astrocytes (Fig. 1h), where a number of genes related to mitochondrial beta-oxidation showed lower expression in the presence of E4 (Supplemental Fig. 3 and 4).

**Fig. 1.**
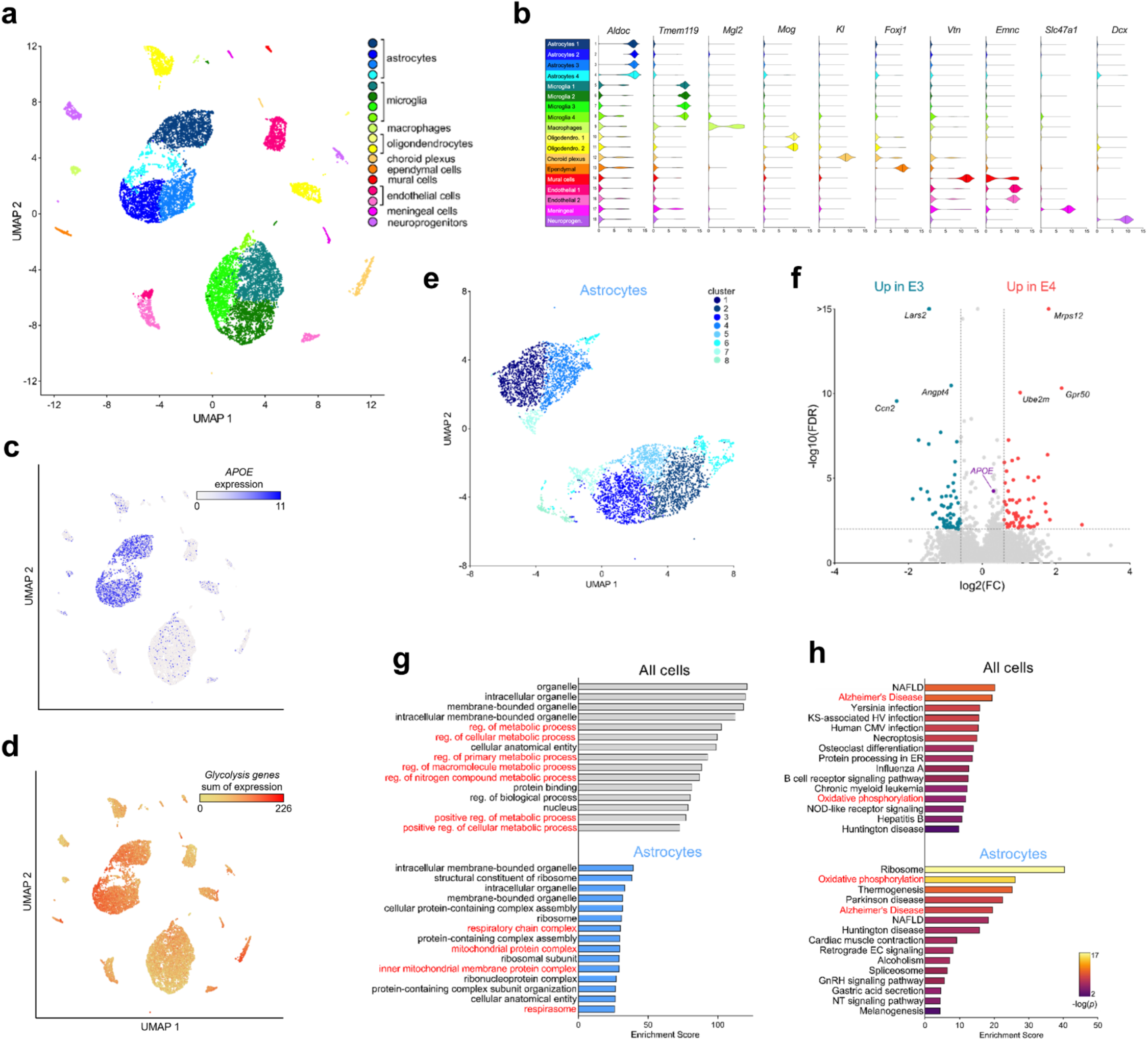
Single-cell RNA sequencing highlights E4-associated changes in glycolysis and oxidative phosphorylation in astrocytes. Whole brain tissue from E3 and E4 mice was digested and subjected to single-cell RNA sequencing (scRNA-seq). **(a)** UMAP visualization of cells from E3 and E4 mouse brains (3 pooled hemi-brains per genotype). Cells are colored by cell type. **(b)** Assignment of clusters to specific cell types based on expression of known gene markers (astrocytes, *Aldoc*; microglia, *Tmem119*; macrophages, *Mgl2*; oligodendrocytes, *Mog*; choroid plexus, *Kl*; ependymal cells, *Foxj1*; mural cells, *Vtn*; Ednothelial cells, *Emnc*; meningeal, *Slc47a1*; neuroprogenitor cells, *Dcx*). **(c-d)** Expression of both *APOE* **(c)** and glycolysis genes **(d)** was highest in astrocyte cell populations. Glycolysis gene expression is shown as the sum of the expression of 39 detected genes belonging to the KEGG pathway “glycolysis and gluconeogenesis”. **(e)** UMAP visualization of astrocytes (Aldoc+ cells). Cells are colored by cluster. **(f)** Volcano plot showing differentially expressed genes in E3 and E4 astrocytes. **(g-h)** Gene ontology **(g)** and pathway enrichment **(h)** analyses highlights APOE-associated gene expression changes in metabolic pathways, particularly mitochondrial complex and oxidative phosphorylation (highlighted in red). *Abbreviations: CMV, cytomegalovirus; EC, endocannabinoid; ER, endoplasmic reticulum; GnRH, Gonadotropin-releasing hormone; HV, herpesvirus; KS, Kaposi sarcoma; NAFLD, non-alcoholic fatty liver disease; NT, Neurotrophin; reg., regulation*.

### Stable isotope resolved metabolomics reveals increased lactate synthesis and decreased glucose entry into the TCA cycle in E4 brains and astrocytes

The single-cell gene expression patterns suggested astrocyte-directed changes in glycolysis and oxidative phosphorylation in E4 cells. Astrocytes are the primary source of both cerebral lactate (the end product of glycolysis) ^45^ and apoE ^38^. Therefore, we next utilized stable isotope resolved metabolomics (SIRM) to quantitatively assess glucose utilization *in vivo* in mice expressing human E3 or E4 and *in vitro* in primary astrocytes expressing human APOE (Fig. 2a). Fasted E3 and E4 mice were administered an oral gavage of fully labeled [U-^13^C] glucose and brain tissue was collected 45 minutes later for mass spectrometry analysis of ^13^C enrichment in central carbon metabolites. Notably, the brains of E4 mice showed significantly higher ^13^C-lactate (fully labeled, m+3) compared to E3 mice (Fig. 3b).

**Fig. 2.**
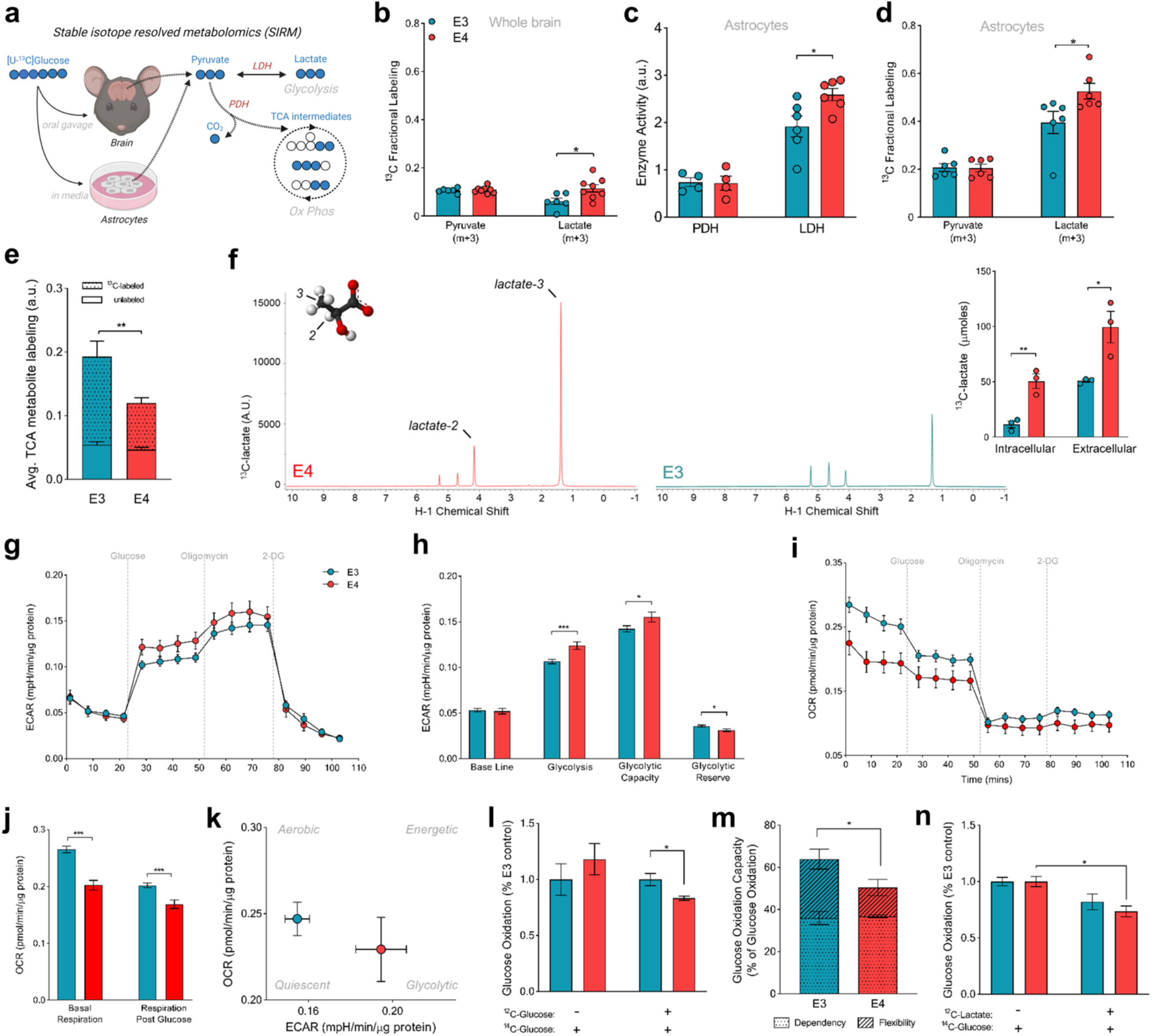
E4 increases lactate production in mouse brain and E4 astrocytes show increased glycolytic flux and lower oxidative respiration. **(a)** Experimental design (^13^C, blue filled circles; ^12^C, white circles; (m+n, where n is the number of ^13^C labeled carbons within a metabolite). [U-^13^C] glucose was administered *in vivo* to E3 (n=6) and E4 (n=8) mice via oral gavage, brain tissue was collected after 45 minutes, and metabolites analyzed for ^13^C enrichment in pyruvate and lactate. E3 and E4 expressing astrocytes were cultured in [U-^13^C] glucose media for 24 hours, media collected, cells washed, and metabolites analyzed for ^13^C enrichment (*n*=6). **(b)** While fully labeled pyruvate is present in similar amounts in E3 and E4 brains, lactate synthesized from ^13^C-glucose is higher in E4 mouse brains. **(c-e)** Primary astrocytes expressing E4 show increased ^13^C enrichment in lactate **(c)**, higher LDH activity **(d)**, and decreased ^13^C enrichment in the TCA cycle (average of all detected TCA intermediates) **(e)**. **(f)** Increased lactate synthesis as measured by HSQCAD NMR spectroscopy (*n*=3). Representative NMR spectra **(f)** showing E4 astrocytes have both increased intracellular ^13^C-lactate and export more lactate into extracellular media **(bar graph insert)**. **(g)** Extracellular acidification rate (ECAR) of E3 and E4 primary astrocytes shown over time during the glycolysis stress test (n=24 for both groups). **(h)** Contributions to ECAR at baseline, in response to glucose (glycolysis), in response to stress (glycolytic capacity), and un-tapped reserve were calculated. **(i)** Oxygen consumption rate (OCR) during the glycolysis stress test assay was graphed over time and **(j)** represented as average respiration before and after glucose. **(k)** Metabolic phenotypes of E3 and E4 astrocytes were characterized by plotting ECAR vs. OCR. **(l)** E3 and E4 astrocytes were incubated in glucose free media (−) or glucose rich media (+) and oxidation of 1.0 μCi/mL ^14^C-glucose was measured by trapping ^14^CO2 and counting radio activity. (*P<0.05 unpaired t-test, two-tailed, *n*=4 per genotype). **(m)** Glucose oxidation capacity, dependency, and flexibility was assessed in E3 and E4 astrocytes via the Mito Fuel Flex Assay. **(n)** E3 and E4 astrocytes were incubated in 1.0 μCi/mL ^14^C-glucose with (+) or without (−) 12.5mM lactate (*n*=3). (b-l,n, *P<0.05, ***P<0.001, ****P<0.0001, unpaired t-test, two tailed) (m, *P<0.05 Two-way ANOVA, Sidak’s multiple comparisons test).

We next incubated primary astrocytes expressing E3 or E4 with [U-^13^C] glucose and collected cell lysates 24 hours later for ^13^C enrichment analysis. E4 astrocytes showed a significant increase in activity of lactate dehydrogenase (LDH) – the enzyme responsible for interconversion of pyruvate and lactate – and in ^13^C-glucose conversion to lactate (Fig. 4c-d). Perhaps unsurprisingly, lactate generation was higher in this glycolytic cell type compared to whole brain homogenates (Fig. 4b vs 4d). Conversely, E4 astrocytes displayed substantially lower ^13^C enrichment of TCA intermediates, suggesting decreased glucose entry into the TCA cycle (Fig. 4e). To confirm these results, we performed an independent ^13^C-glucose tracing experiment in immortalized astrocytes expressing human E3 or E4 ^46^ and quantified ^13^C-lactate production using nuclear magnetic resonance (NMR) spectroscopy (Fig. 4f). Again, E4 astrocytes showed significantly higher lactate synthesis, as evidenced by increased ^13^C-lactate both intracellularly and in the media (Fig. 4f, insert). Together, these data describe an E4-associated increase in glucose flux into late glycolysis at the expense of entry into the TCA cycle for oxidative phosphorylation.

### E4 astrocytes exhibit impairments in glucose oxidation

To functionally assess astrocyte glycolytic flux *in vitro,* we measured the extracellular acidification rate (ECAR, a marker of glycolysis and lactate export) before and after glucose injection. E4 astrocytes displayed significantly higher ECAR after addition of glucose compared to E3 astrocytes, as well as a higher glycolytic capacity, suggesting these cells shunt more glucose to lactate (Fig. 2g-h). E4 astrocytes also displayed a significantly lower oxygen consumption rate (OCR), both before and after addition of glucose to the media, suggesting an inherent reduction in oxidative metabolism (Fig. 2i-j). Together these data further support an E4-associated shift toward glycolysis (Fig. 2k). We next measured glucose oxidation by treating astrocytes with radiolabeled ^14^C-glucose and capturing the oxidative product ^14^CO_2_. E4 astrocytes oxidized less glucose to CO_2_ compared to E3, but only when the radiolabel ([nM]) was given with a substantial amount of non-labeled glucose ([mM]) (Fig. 2l). E4 astrocytes also displayed decreased capacity and flexibility in regards to glucose oxidation, as they were relatively unable to increase glucose oxidation when other fuel sources (fatty acids and glutamine) were inhibited (Fig. 2m). We reasoned that lower rates of glucose oxidation in a glucose rich environment in E4 cells may be due to increased conversion of glucose to lactate, which in turn inhibits downstream oxidative processes ^47^. Therefore, we tested glucose oxidation following lactate supplementation, and found that E4 astrocytes oxidize less glucose in the presence of lactate than E3 astrocytes (Fig. 2n). Together, these results suggest that E4 astrocytes exhibit increased reliance on aerobic glycolysis and are less flexible and less able to oxidize glucose, a phenotype seemingly exacerbated by a high glucose environment and/or the presence of lactate.

### E4 mice fail to increase energy expenditure on a high carbohydrate diet

Given the apparent shift toward aerobic glycolysis in the brain and astrocytes of mice expressing APOE4, we next asked if this metabolic reprogramming was a global phenomenon (i.e. could it be detected with whole body measures). Indirect calorimetry (IC) assesses energy expenditure by measuring metabolic gases to calculate the energy released when substrates are oxidized. Energy expenditure (EE) is estimated using the Weir equation (EE = 3.9 * VO_2_ + 1.11 VCO_2_), with the assumption that anaerobic respiration is negligible and substrates are fully oxidized to CO_2_ ^24^. However, this assumption is confounded when energy is derived through non-oxidative processes such as aerobic glycolysis – a phenomena in which glucose is fully metabolized to lactate despite normoxia ^48^. To test whether mice expressing APOE4 display an aerobic glycolysis related shift in metabolism, we used IC to track energy expenditure in mice expressing human E3 or E4. Young mice carrying the human E4 allele exhibited significantly lower EE, VCO_2_, and VO_2_ compared to young E3 mice during their inactive period (light cycle) (Fig. 3a-c). Since mouse IC cages allow for a prolonged and controlled assessment of metabolism, we provided a long-term glucose challenge by way of a high carbohydrate diet (HCD). Interestingly, these E4-associated decreases were exaggerated following introduction of the HCD; E4 mice again showed substantially lower EE, VCO_2_, and VO_2_ compared to E3 mice (Fig. 3d-f). Further, when we analyzed the HCD-induced change in EE, VCO_2_, and VO_2_ from baseline (normal chow), we found both genotypes to show significant positive changes except for E4 VO_2_ (Fig 3i). This suggests that E4 mice fail to increase oxygen consumption in response to excess dietary carbohydrates. These changes occurred independently of differences in activity and food intake, and was not explained by differences in body weight (Fig. j-l). Together, these data suggest that E4 acts in young mice to lower energy expenditure via a mechanism outside of the typical contributions of feeding, body mass, and activity.

**Fig. 3.**
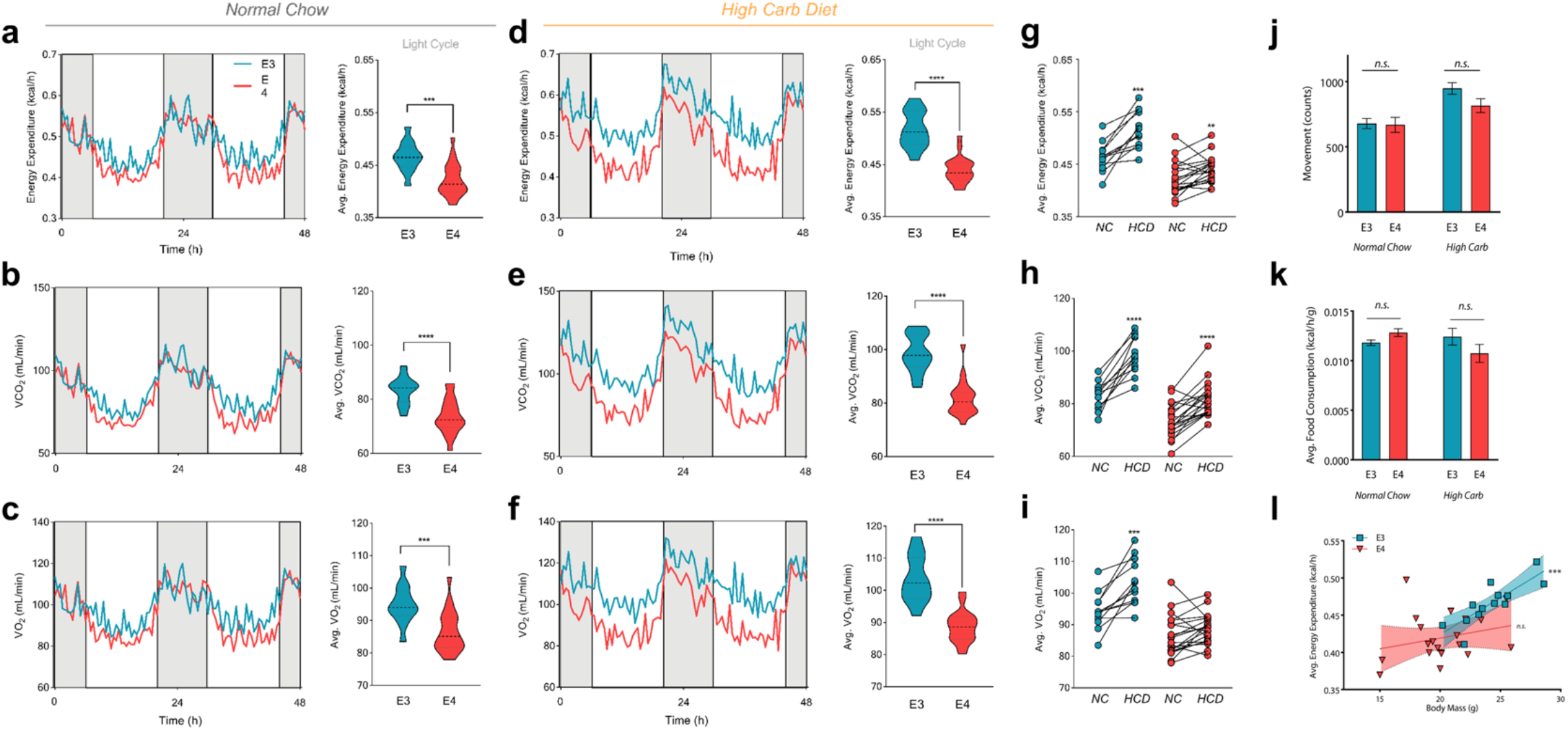
E4 mice have lower energy expenditure and fail to increase oxygen consumption following a high carbohydrate diet. **(a-f)** E3 and E4 mice were housed individually for 48 hours with *ad libitum* access to normal chow **(a-c)** or a high carbohydrate diet (HCD) **(d-f)** and energy expenditure (EE), VCO_2_ and VO_2_ were measured. Dark cycles are indicated in grey with light cycles in white. Light cycles were used for calculating averages of EE, VCO_2_ and VO_2_ (shown to the right) (\*\*\**P*<0.0001, \*\*\*\**P*<0.00001, unpaired *t*-test, two-tailed; E3 *n*=13, E4 *n* = 20). **(j)** Activity and **(k)** food consumption during light cycles were averaged for E3 and E4 mice (E3 *n*=13, E4 *n* = 20). **(l)** Analysis of covariance was performed by separately correlating average EE and body weight for E3 and E4 mice. (Spearman correlation r=0.86, ***P<0.001).

### Young female E4 carriers have a lower resting energy expenditure

We next asked if this E4-associated shift toward aerobic glycolysis observed in cell and animal models translated to APOE4+ humans. To test this, we used IC to test the effect of *APOE* on whole body metabolism in a cohort of healthy, cognitively normal young and middle-aged volunteers (Supplemental Tables 2 and 3). Using a mobile metabolic cart designed to measure VO_2_ and VCO_2_, we assessed exhaled breath measures of volunteers at rest, during a cognitive task, and after a glucose challenge (Fig. 4a and Supplemental Fig. 5). We began each session by assessing the resting energy expenditure (REE) and respiratory exchange ratio (RER) of participants. After a five-minute buffer to achieve steady state ^26,27^, we recorded REE over a 25 minute period at 15 second intervals and averaged the RER and REE for each individual. There was no *APOE* effect on RER (Supplemental Fig. 6). Consistent with previous studies, we found REE and age to be negatively correlated (Fig. 4c). However, when we stratified our analysis by E4 status, linear regression revealed significantly different slopes between carriers and non-carriers, suggesting an E4-associated confound in the age versus energy expenditure relationship (Fig. 4d).

**Fig. 4.**
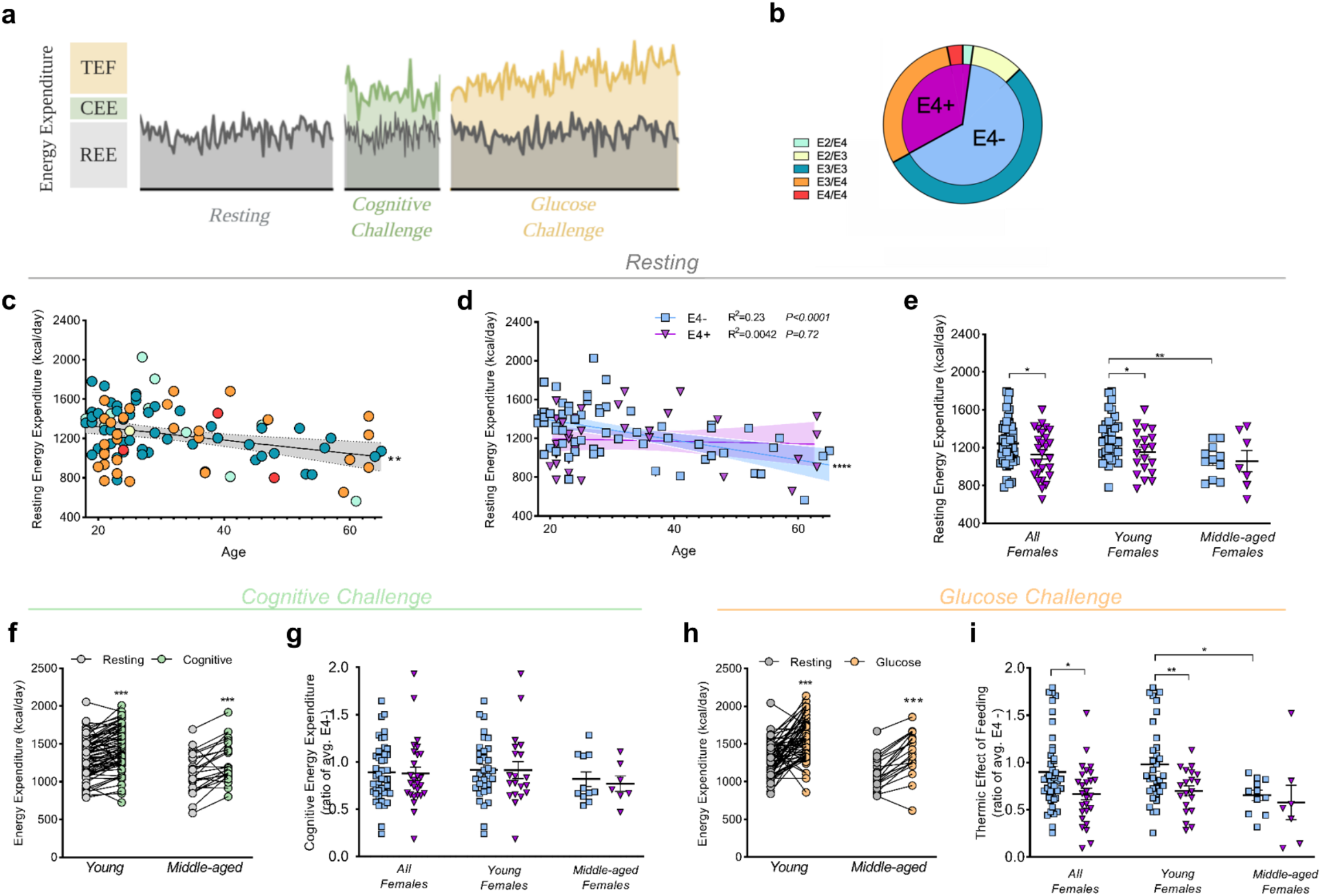
Female E4 carriers show lower resting energy expenditure and lower thermic effect of feeding after a glucose challenge. **(a)** Experimental design of study. Individual components of energy expenditure (EE) were assessed in three distinct periods. Resting energy expenditure (REE) was assessed during the resting period. Cognitive energy expenditure (CEE) was assessed during the cognitive challenge and defined as difference in the area under the curve (AUC) of EE during the cognitive challenge and the AUC of EE from the resting period. Thermic effect of feeding (TEF) was assessed during the glucose challenge and calculated as the difference in AUC of EE during the glucose challenge and AUC of REE. **(b)** *APOE* genotypes of subjects represented in the study (E4-*n*=61, E4+ *n*=33; E2/E4 *n*=2, E2/E3 *n*=10, E3/E3 *n*=51, E3/E4 *n*=28, E4/E4 n=3). **(c)** Correlation of average REE with participant age (Pearson correlation R^2^=0.11, \*\**P*<0.01, *n*=94). **(d)** Correlation of average REE and participant age separated by E4 carriers (purple) and non-carriers (blue) (E4-R^2^=0.233, \*\*\*\**P*<0.0001; E4+ R^2^ =0.0042, *P*=0.719, E4-*n*=61 and E4+ *n*=33). Shaded areas refer to 95% confidence intervals. **(e)** Average REE for all, young, and middle-age E4-(*n*=44, 33, and 11 respectively) and E4+ females (*n*=27, 20, and 7 respectively) (\**P*<0.05, \*\**P*<0.01, unpaired *t*-test, two-tailed). **(f)** Average EE between resting and cognitive test periods in young (*n*=71) and middle-aged (*n*=23) participants. (\*\*\**P*<0.001, paired *t*-test, two-tailed). **(g)** CEE for all female participants and for the two age cohorts. **(h)** Average EE between resting and glucose challenge periods in young and middle-aged participants (\*\*\**P*<0.001, paired *t*-test, two-tailed). **(i)** TEF for all females and for the two age cohorts, further separated by E4 carriers and non-carriers. (\**P*<0.05, unpaired *t*-test, two-tailed).

We then separated E4+ and E4- individuals into young (<40 years of age) and middle aged (40-65 years of age) cohorts based on previous literature ^4,17^. After adjusting for covariates, we observed a significantly lower REE in female E4 carriers compared to non-carriers, particularly in the young cohort (Fig. 4e). This E4 effect on REE was not significant in males (Supplemental Fig. 7), together suggesting that there is no age-related REE decline in E4 carriers, and that the energy expenditure-*APOE* interaction is modified by sex, with female E4 carriers displaying lower REE.

### E4 does not alter cognitive energy expenditure

Given the critical role of *APOE* in modulating cognitive function and dementia risk, we next tested if a mental stressor would reveal further genotype-specific differences in energy expenditure. To avoid potential confounding readouts of movement, subjects were asked to remain perfectly still while completing a challenging Novel Image Novel Location test (Supplemental Fig. 5c). We observed a significant increase in average EE during the cognitive challenge in all subjects (Fig. 4f). However, we found no difference in cognitive energy expenditure (CEE), nor in test response accuracy, between E4 carriers and non-carriers (Fig. 4g and Supplemental Fig. 8). To our knowledge, only two other studies have attempted to utilize IC to quantify the contribution of cerebral activation (i.e. a mental task) to whole body metabolic measures ^49,50^. While we did not observe an *APOE* effect on metabolic measures during the cognitive challenge, we did find that IC is a sensitive tool to evaluate metabolic changes due to mental stress, as all participants showed a significant increase in energy expenditure (Fig. 4f).

### Female E4 carriers have a blunted increase in oxygen consumption after a dietary glucose challenge

We next sought to measure the thermic effect of feeding (TEF) - a constituent of EE that indicates the energy used to absorb, digest, and metabolize dietary energy ^51,52^. To induce TEF, all participants consumed a high carbohydrate drink in less than two minutes (Supplemental Fig. 5d). Energy expenditure during the dietary challenge increased significantly in all participants (Fig. 4h), and similar to resting EE, young female E4 carriers displayed a significantly lower TEF than non-carriers (Fig. 4i).

Plotting the time course of EE after participants consumed the glucose drink revealed a dramatically blunted energy response in E4+ individuals, an effect driven by E4+ females (Fig. 5a and Supplemental Fig. 9). Further stratification by individual genotypes showed a clear stepwise effect of *APOE* (Fig. 5b). Post-glucose drink VCO_2_ values revealed a similar, but non-significant trend of lower CO_2_ production in E4 carriers (Supplemental Fig. 9). Importantly, we observed that while non-carriers significantly increased their oxygen consumption following the glucose drink, female E4 carriers did not, as noted by significant E4-associated decreases in total oxygen consumption across the post-glucose period (Fig. 5c-d).

**Fig. 5.**
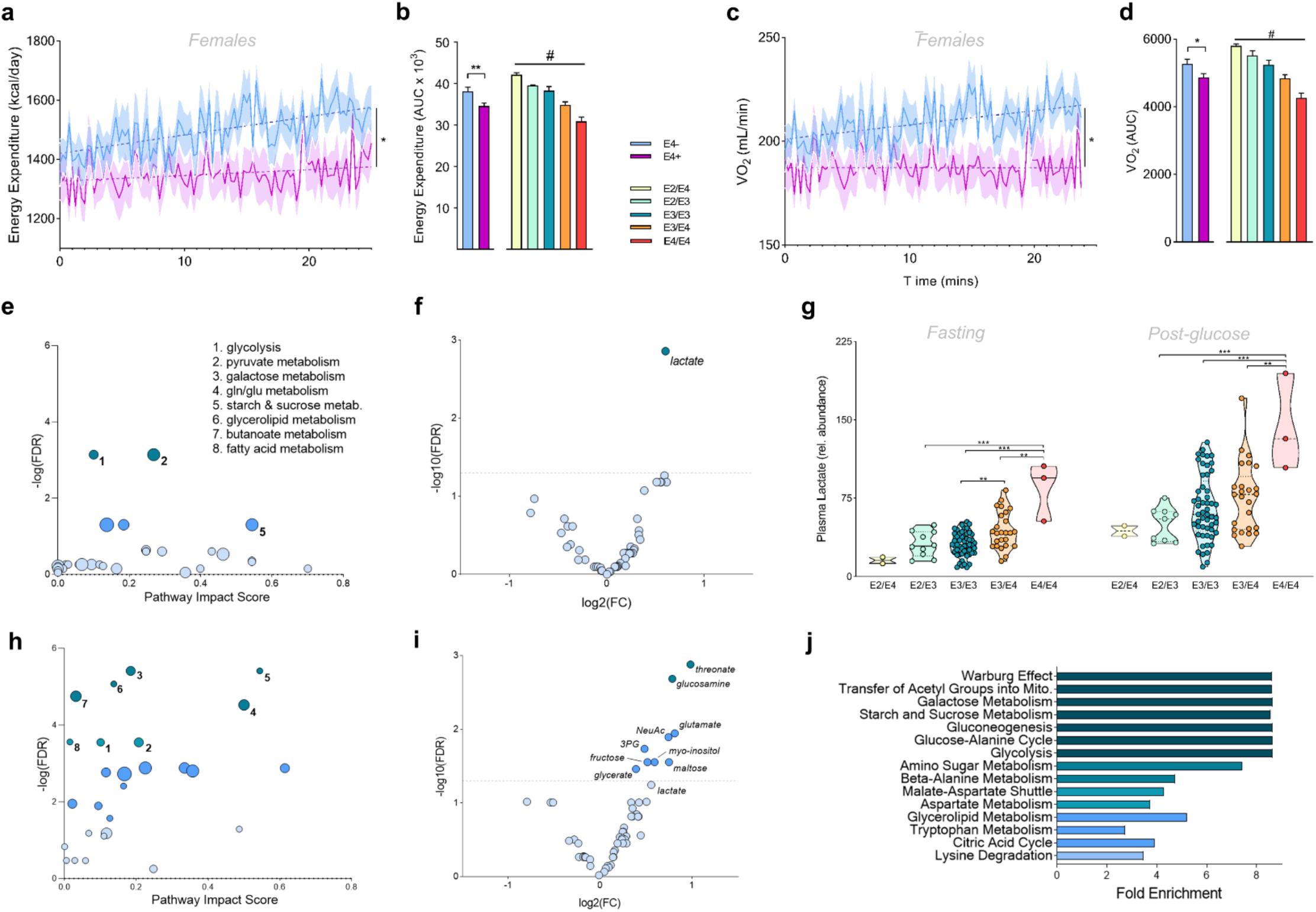
E4 carriers show lower energy expenditure, decreased oxygen consumption, and pro-glycolytic changes in the plasma metabolome. **(a,c)** Energy expenditure (EE) **(a)** and VO_2_ **(c)** of female E4 carriers (purple) and E4 non-carriers (blue) during the glucose challenge. Values shown are means (lines) +/− SEM (shaded). (E4-*n*=44, E4+ *n*=27; \**P*<0.05, Two-way ANOVA repeated measures) **(b,d)** Incremental area under the curve (AUC) of EE **(b)** and VO_2_ **(d)** was determined by E4 carriage and further by respective *APOE* genotypes in all participants. (E4-*n*=61, E4+ *n*=33; E2/E4 *n*=2, E2/E3 *n*=10, E3/E3 *n*=51, E3/E4 *n*=28, E4/E4 *n*=3) (\**P*<0.05, \*\**P*<0.01, unpaired *t*-test, two-tailed; #*P*<0.05 One-way ANOVA). (**e,h**) Pathway impact analysis highlights pyruvate metabolism and glycolysis as pathways significantly altered by E4 carriage in human plasma at baseline **(e)**, while multiple carbohydrate and lipid processing pathways are altered by E4 carriage following the glucose drink **(h)** (FDR <0.01). **(f,i)** Volcano plots showing changes in plasma metabolites. Lactate was the most significantly altered metabolite by *APOE* genotype at baseline **(f)**, while multiple metabolites differed post-glucose drink **(i)**(ANOVA, FDR <0.05). **(g)** Lactate values in individual subjects as determined by GC-MS analysis. **(j)** Enrichment analysis highlights multiple metabolic pathways as significantly altered by E4 carriage following the glucose drink, including the top hit of ‘Warburg effect’. All comparisons are E4+ (*n*=33) vs E4- (*n*=61).

### Targeted metabolomics reveals glycolysis as a differentially regulated pathway in E4+ plasma

To determine if the observed *APOE* differences in energy expenditure were reflected in the plasma metabolome, we conducted a targeted metabolomics analysis of human plasma samples before and after the glucose challenge (Extended Data Table 4). A pathway analysis of the plasma metabolome before the glucose drink highlighted E4-associated differences in glycolysis and pyruvate metabolism (Fig. 5e), and further analyses of individual metabolites revealed lactate as the metabolite most strongly affected by E4 carriage (Fig. 5f). Indeed, E4 carriers displayed dramatically higher plasma lactate concentrations before and after the glucose drink (Fig. 5g, Extended Data Fig. 11). Following the glucose challenge, there was an increase in the number of carbohydrate processing pathways and metabolites that were differentially altered in E4 carriers (Fig. 5h-i), and a pathway enrichment analysis highlighted top hits of “Warburg effect” and “Transfer of acetyl groups into mitochondria” (Fig. 5j). Together, analysis of the plasma metabolome from cognitively E4+ individuals suggests a preference for aerobic glycolysis compared to non-carriers.

## Discussion

In the current study, we used indirect calorimetry to show that APOE4 reduces energy expenditure in a cohort of young, cognitively normal females, a phenomenon exacerbated by a dietary glucose challenge. Analysis of the plasma metabolome revealed E4-associated increases in pathways related to carbohydrate processing, specifically aerobic glycolysis, highlighted by higher concentrations of the glycolytic end-product lactate. By applying single-cell RNA sequencing and stable isotope-resolved metabolomics *in vivo,* along with functional assays of cellular respiration *in vitro*, we discovered that both E4 expressing mouse brains and E4 expressing astrocytes increase glucose flux through aerobic glycolysis at the expense of TCA cycle entry and oxidative phosphorylation. Cumulatively, these data highlight a novel mechanism whereby E4 lowers energy expenditure in young women and decreases glucose oxidation by redirecting flux through aerobic glycolysis.

These results are congruent with other studies of APOE4 and AD. For example, a recent study by our group demonstrated that E4 astrocytes have increased lactate production ^53^, and neurons expressing E4 exhibit increased reliance on glycolysis for ATP production with apparent deficits in mitochondrial respiration ^54^. Interestingly, another study showed that fibroblasts from AD patients show a ‘Warburg-type’ (aerobic glycolysis) shift from oxidative phosphorylation to glycolysis with increased lactate production ^55^. Aerobic glycolysis refers to the metabolism of glucose to lactate instead of the oxidative TCA cycle, despite the presence of abundant oxygen. In the brain, this phenomenon occurs in young individuals with a peak around five years of age (when 30% of cerebral glucose is processed anaerobically), and then steadily declines with age ^56^. Aerobic glycolysis in the brain appears to be cell and region specific, with astrocytes playing a major role in certain regions such as the dorsolateral prefrontal cortex, precuneus, and the posterior cingulate cortex ^57^. Importantly, areas associated with aerobic glycolysis also overlap with areas known to accumulate amyloid β, indicating that the anaerobic metabolism of certain brain regions may possibly predict amyloid burden in later life ^58^. Furthermore, recent proteomic profiling of over 2,000 AD brain samples revealed that changes in the expression of proteins involved in glial metabolism was the most significant module associated with AD pathology and cognitive decline ^59^. Increased expression of enzymes in this module included lactate dehydrogenase, pyruvate kinase, and glyceraldehyde-3-phosphate dehydrogenase, all of which are elevated in aerobic glycolysis phenotypes.

Interestingly, recent evidence has shown that lactate is an energy substrate used by the brain ^60^ and a competitive glucose alternative ^61–63^. Lactate has also been shown to decrease FDG-PET signal ^64^. An increase in astrocyte-derived lactate in E4 carriers may compete with glucose as a substrate for brain metabolism and decrease CMRglc. Further, an increase in aerobic glycolysis might also act to lower energy expenditure, as glycolysis produces only 2 moles of ATP compared to the 34 moles of ATP from a mole of glucose metabolized via mitochondrial oxidative phosphorylation. This balance of anaerobic glycolysis versus oxidative phosphorylation behaves reciprocally ^65^. Increased mitochondrial ATP production downregulates glycolysis, while glycolytic ATP synthesis can suppress aerobic respiration ^66^. Given our findings of lower O_2_ consumption and increased production of lactate, we speculate that E4 carriers have lower energy expenditure due to glycolysis being less energetically costly than downstream pathways.

Our study has several limitations. As a primary goal was to assess individual metabolic responses to glucose, we performed blood draws immediately prior and immediately after the glucose challenge. It may be possible that mental stress during the cognitive challenge (which occurred prior to the first blood draw) altered the plasma metabolome beyond the normal resting state. Another potential confounder is that we provided glucose in the form of a sugary milk drink for the glucose challenge. While we used 50g of sugar based on clinical guidelines for glucose challenges ^67^, milk also includes fats and proteins. However, the high relative content of carbohydrates to other macronutrients ensures that any observed response (particularly at the ~30 minute time point analyzed) can be primarily attributed to carbohydrate metabolism. Indeed, the pathways most altered by the glucose challenge included galactose metabolism, starch and sucrose metabolism, and glycolysis. Finally, while we found E2 to be associated with lower plasma lactate and higher EE relative to non-E2 carriers, the study did not include any homozygous E2 carriers and the low overall allele frequency makes interpretation challenging. Still, these results are intriguing based on E2 being a known protective allele for AD ^2,4^, and further study of energy expenditure and glucose metabolism in E2 carriers is warranted.

Current understanding of the development of late-onset AD supports a triad of primary risk factors: E4, female sex, and old age. However, detecting symptoms of eventual cognitive decline in young asymptomatic individuals is critical for primary prevention of AD ^68^. Given the largely disappointing trial outcomes of drugs targeting AD neuropathology ^69^, these therapies may be intervening after a ‘point of no return’ and thus offer minimal benefit in prognosis ^70^. In order to design therapies for early interventions in those at risk for AD, we must first identify measurable biomarkers whose severity and/or change over time correlate with risk for clinically observable AD. In the current study, we used indirect calorimetry to show that *APOE4* reduces energy expenditure in a cohort of young cognitively normal females, a phenomenon exacerbated by a dietary glucose challenge. While using indirect calorimetry for metabolic studies is common in clinical settings and exercise studies ^71,72^, to our knowledge the method has not been previously applied to investigate biomarkers of cognitive impairment. Thus, repurposing IC to study the metabolic effects of an AD risk factor such as E4 represents a mobile, simple, and cost-effective new approach.

Although resting energy expenditure was significantly lower in female E4 carriers at rest, the most striking effect of *APOE* was observed after participants underwent a dietary carbohydrate challenge. There, E4+ individuals failed to increase VO_2_, leading to a significantly lower EE compared to non-carriers. Energy expenditure is estimated using the Weir equation (EE = 3.9 * VO_2_ + 1.11 VCO_2_), with the assumption that anaerobic respiration is negligible and substrates are fully oxidized to CO_2_ ^24^. However, this assumption is confounded when energy is derived through non-oxidative processes such as aerobic glycolysis – a phenomena in which glucose is fully metabolized to lactate despite normoxia ^48^. Given the decreased VO_2_ and increased plasma lactate concentrations in E4+ subjects, we hypothesize that these individuals are diverting a higher fraction of glucose to aerobic glycolysis as opposed to oxidative phosphorylation. Along these lines, analysis of the plasma metabolome revealed E4-associated increases in pathways primarily related to carbohydrate processing, specifically aerobic glycolysis. These results were in line with our results from mouse and cell models of APOE4, where our application of scRNAseq, stable isotope-resolved metabolomics and functional assays of cellular respiration showed that both E4 expressing mouse brains and E4 expressing astrocytes increase glucose flux through aerobic glycolysis at the expense of TCA cycle entry and oxidative phosphorylation.

Cumulatively, these data highlight a novel mechanism whereby E4 lowers energy expenditure and decreases glucose oxidation by redirecting flux through aerobic glycolysis. While many questions remain, our study highlights novel roles for *APOE* and sex in modulating systemic and cerebral glucose metabolism and provides a feasible method to assess *APOE*-dependent metabolic signatures in pre-symptomatic young individuals. These findings provide important insights that may help to define dietary and pharmacological approaches to delay or prevent incipient AD in high-risk individuals.

## Acknowledgments

The authors thank the Center for Clinical and Translational Science nursing staff for assistance with venipunctures, Dr. Arnold Stromberg, Dr. Richard Kryscio and Ning Li for statistics consultation, and Anna Wilwerding for her assistance in data input and organization.

## Funding

This work was supported by the National Institute on Aging (BCF - F30AG06342201A; HCW-1T32AG057461-01; JMM - 1RF1NS118558-01; LAJ - 1R01AG060056 and R01AG062550), the National Institute of General Medical Sciences (HCW-5T32GM118292-03, LAJ-COBRE P20 GM103527, RCS - COBRE P20 GM121327), the National Institute of Neurological Disorders and Stroke (MSG - R01NS070899, R01NS070899-09S2, R35NS116824), the American Cancer Society Institutional Research Grant (RCS -#16-182-28), and the St Baldrick’s Foundation (Career Development Award to RCS). The project described was also supported by the National Center for Advancing Translational Sciences, National Institutes of Health (UL1TR001998) and the Redox Metabolism Shared Resource Facility of the University of Kentucky Markey Cancer Center (P30CA177558).

## Author contributions

BCF helped design the study, performed the experiments, and wrote the paper. HCW, MAP, and LEAY performed the plasma metabolomics analysis and stable isotope-resolved metabolomics experiments. NAD and JMM performed scRNAseq experiments and analyzed data. DJC performed mouse indirect calorimetry experiments and assisted with human subject recruitment and testing. GKN, DJC, and RK assisted in the recruitment and testing of human subjects. RM, JCK and GH assisted with data analysis and *APOE* genotyping. JAB performed in vitro measures of cell metabolism. AEW and EJA assisted with the plasma metabolomics data curation and analysis. VAG served as the medical monitor for the study. PAK guided experimental design and provided assistance and expertise in the testing of human subjects. MSG and RCS provided resources, expertise and guidance to complete the stable isotope resolved metabolomics experiments. LAJ conceived and designed the study, analyzed and interpreted data, and wrote the paper. All authors read the manuscript and provided revisions.

## Competing interests

The authors have no conflicts of interest.

## Data and materials availability

Clinical Trial #: NCT03109661

## Supplemental Material

**Supplemental Table 1.**
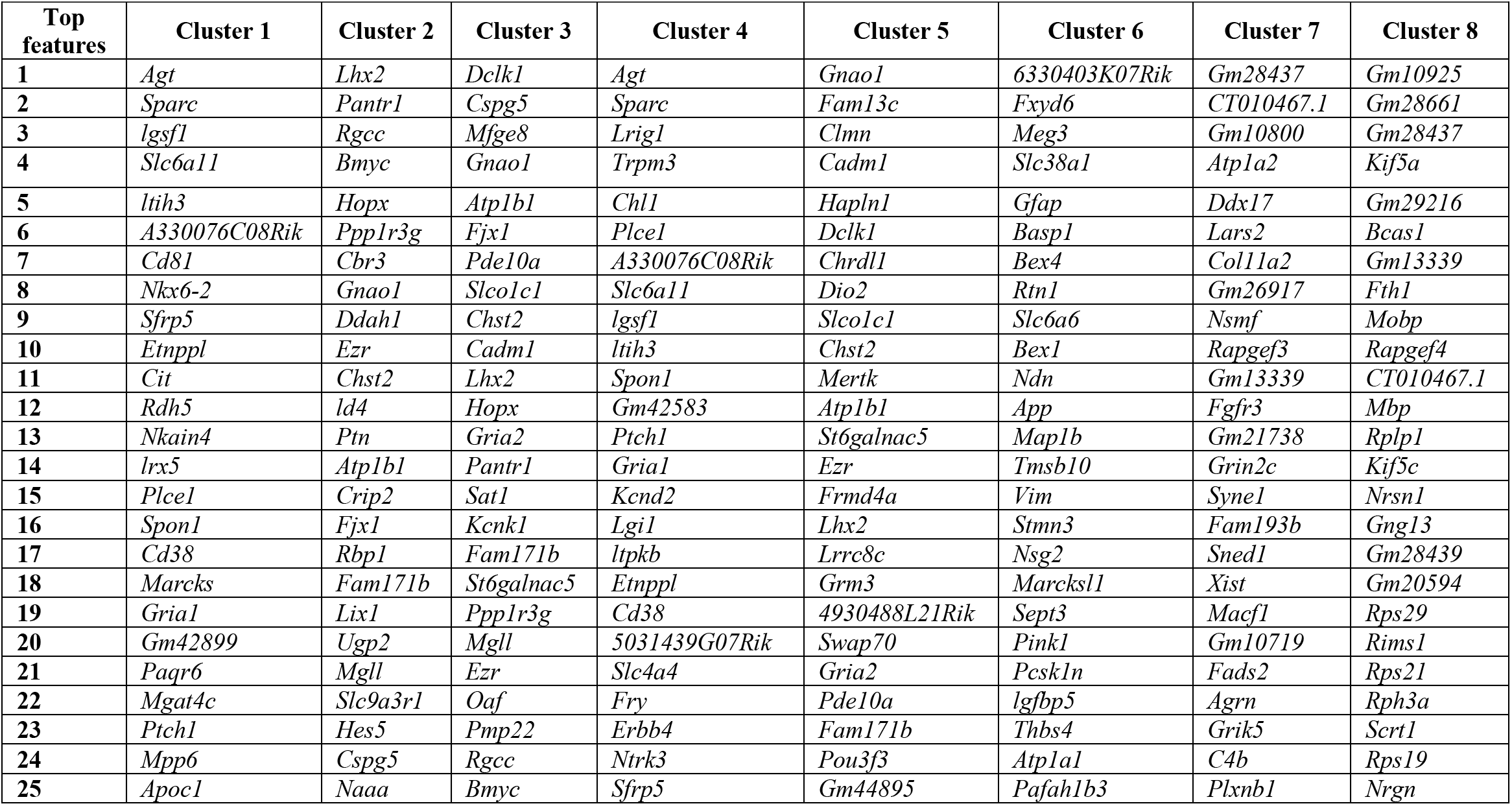
Top 25 genes classifying the eight distinct astrocyte clusters identified in scRNAseq analysis of E3 and E4 mouse brains (clusters visualized in Fig. 4)

**Supplemental Table 2.** Age, sex, and *APOE* genotype of cognitively normal individuals according to E4 carriage and age cohort (young=18-39, middle-aged=40-65). Values represent means +/− (SD).

|  | Age |  | Sex (N) |  | APOE Genotype (N) |  |  |  |  |
| --- | --- | --- | --- | --- | --- | --- | --- | --- | --- |
|  | Average Age (SD) | P | Male | Female | APOE 2/3 | APOE 3/3 | APOE 2/4 | APOE 3/4 | APOE 4/4 |
| Young |  |  |  |  |  |  |  |  |  |
| E4 Non-carriers (E4-) | 24.91<br>(4.84) | 0.486 | 13 | 33 | 8 | 38 | 0 | 0 | 0 |
| E4 Carriers (E4+) | 25.84<br>(6.08) |  | 5 | 20 | 0 | 0 | 2 | 21 | 2 |
| Middle-age |  |  |  |  |  |  |  |  |  |
| E4 Non-carriers (E4-) | 51.18<br>(7.79) | 0.233 | 4 | 11 | 2 | 13 | 0 | 0 | 0 |
| E4 Carriers (E4+) | 55.5<br>(8.78) |  | 1 | 7 | 0 | 0 | 0 | 7 | 1 |

**Supplemental Table 3.** Clinical characteristics of cognitively unimpaired individuals according to E4 carriage and age cohort (young=18-39, middle-aged=40-65). Values represent means +/− (SD). Ca, Caucasian; AA, African American; His, Hispanic; A, Asian; BMI, body mass index.

|  | Race (N) |  |  |  | Vitals |  |  |  |  |  |  |  |  |  |
| --- | --- | --- | --- | --- | --- | --- | --- | --- | --- | --- | --- | --- | --- | --- |
|  | Ca | AA | His | A | BMI<br>(kg/m²) | P | Waist/<br>Hip Ratio | P | Systolic<br>(mmHg) | P | Diastolic<br>(mmHg) | P | Lean<br>Body<br>Mass (kg) | P |
| Young |  |  |  |  |  |  |  |  |  |  |  |  |  |  |
| E4- | 34 | 4 | 3 | 4 | 25.15<br>(3.83) | 0.93 | 0.86<br>(0.1) | 0.56 | 123.2<br>(21.2) | 0.79 | 72.3<br>(14.4) | 0.9 | 48.2<br>(7.3) | 0.33 |
| E4+ | 22 | 3 | 0 | 0 | 25.06<br>(4.72) |  | 0.87<br>(0.1) |  | 121.8<br>(18.5) |  | 72.1<br>(12.1) |  | 46.4<br>(7.2) |  |
| Middle-age |  |  |  |  |  |  |  |  |  |  |  |  |  |  |
| E4- | 12 | 3 | 1 | 0 | 25.83<br>-6.07 | 0.45 | 0.9<br>(0.1) | 0.14 | 123.4<br>(17.6) | 0.76 | 74.2<br>(8.7) | 0.6 | 47.1<br>(7.7) | 0.63 |
| E4+ | 7 | 0 | 1 | 0 | 27.92<br>-6.73 |  | 0.81<br>(0.2) |  | 126<br>(23.1) |  | 76.8<br>(15.9) |  | 48.9<br>(9.2) |  |

**Supplemental Table 4.** Plasma metabolites of study participants analyzed by gas chromatography – before and after a dietary glucose challenge.

| *(all comparisons shown are E4+ vs E4-)* |  | Pre-glucose challenge |  | Post-glucose challenge |  |
| --- | --- | --- | --- | --- | --- |
| Metabolite | HMDB ID | Ratio | FDR | Ratio | FDR |
| beta-alanine | HMDB0000056 | 1.059 | 0.797 | 1.179 | 0.314 |
| cholesterol | HMDB0000067 | 1.176 | 0.541 | 1.084 | 0.699 |
| citrate | HMDB0000094 | 1.258 | 0.273 | 1.276 | 0.152 |
| GABA | HMDB0000112 | 1.203 | 0.532 | 1.419 | 0.096 |
| glyoxylic acid | HMDB0000119 | 0.822 | 0.489 | 0.891 | 0.542 |
| glycine | HMDB0000123 | 1.102 | 0.717 | 1.098 | 0.546 |
| glycerol 3-phosphate | HMDB0000126 | 1.002 | 0.994 | 0.853 | 0.542 |
| fumaric acid | HMDB0000134 | 0.940 | 0.797 | 0.822 | 0.314 |
| glyceric acid | HMDB0000139 | 1.243 | 0.193 | 1.310 | **0.034** |
| glutamic acid | HMDB0000148 | 1.427 | 0.066 | 1.757 | **0.011** |
| ethanolamine | HMDB0000149 | 1.141 | 0.532 | 1.160 | 0.291 |
| tyrosine | HMDB0000158 | 1.072 | 0.797 | 1.331 | 0.153 |
| phenylalanine | HMDB0000159 | 0.999 | 0.994 | 1.141 | 0.354 |
| maltose | HMDB0000163 | 1.143 | 0.775 | 1.681 | **0.028** |
| threonine | HMDB0000167 | 0.597 | 0.107 | 0.897 | 0.546 |
| isoleucine | HMDB0000172 | 1.063 | 0.798 | 1.183 | 0.299 |
| lysine | HMDB0000182 | 0.929 | 0.797 | 1.197 | 0.216 |
| lactose | HMDB0000186 | 1.218 | 0.466 | 1.471 | 0.057 |
| serine | HMDB0000187 | 1.025 | 0.919 | 1.277 | 0.118 |
| lactate | HMDB0000190 | 1.519 | **0.001** | 1.264 | **0.013** |
| oleic acid | HMDB0000207 | 0.722 | 0.294 | 0.991 | 0.957 |
| α-ketoglutarate | HMDB0000208 | 1.084 | 0.775 | 1.057 | 0.715 |
| myo-inositol | HMDB0000211 | 1.149 | 0.532 | 1.508 | **0.028** |
| n-acetylgalactosamine | HMDB0000212 | 1.507 | 0.054 | 1.359 | 0.275 |
| palmitic acid (polar lipids) | HMDB0000220 | 1.074 | 0.797 | 1.341 | 0.098 |
| n-acetyl-neuraminic acid | HMDB0000230 | 1.026 | 0.919 | 1.676 | **0.013** |
| pyruvate | HMDB0000243 | 1.128 | 0.532 | 1.108 | 0.464 |
| sucrose | HMDB0000258 | 0.817 | 0.649 | 0.791 | 0.326 |
| serotonin | HMDB0000259 | 0.757 | 0.242 | 1.032 | 0.861 |
| pyroglutamic acid | HMDB0000267 | 1.256 | 0.312 | 1.336 | 0.153 |
| ribose | HMDB0000283 | 0.9659 | 0.894 | 1.102 | 0.546 |
| urea | HMDB0000294 | 1.1415 | 0.621 | 1.117 | 0.441 |
| creatinine | HMDB0000562 | 0.7535 | 0.445 | 0.701 | 0.098 |
| fructose | HMDB0000660 | 1.212 | 0.489 | 1.429 | **0.028** |
| linoleic acid polar | HMDB0000673 | 0.914 | 0.797 | 1.066 | 0.750 |
| methionine | HMDB0000696 | 1.268 | 0.358 | 1.205 | 0.313 |
| homoserine | HMDB0000719 | 0.857 | 0.647 | 0.688 | 0.098 |
| malic acid | HMDB0000744 | 1.185 | 0.441 | 1.185 | 0.247 |
| 3-phosphoglyceric acid | HMDB0000807 | 1.306 | 0.0843 | 1.398 | **0.018** |
| stearic acid | HMDB0000827 | 1.156 | 0.532 | 1.317 | 0.143 |
| valine | HMDB0000883 | 1.174 | 0.533 | 1.217 | 0.284 |
| tryptophan | HMDB0000929 | 0.957 | 0.822 | 1.105 | 0.54 |
| threonic acid | HMDB0000943 | 1.464 | 0.065 | 1.977 | **0.0013** |
| F16BP | HMDB0001058 | 0.783 | 0.242 | 0.927 | 0.584 |
| glucosamine-1-phosphate | HMDB0001109 | 0.739 | 0.193 | 0.883 | 0.527 |
| alanine | HMDB0001310 | 1.018 | 0.919 | 1.192 | 0.244 |
| guanosine-5-phosphate | HMDB0001397 | 1.063 | 0.797 | 0.848 | 0.353 |
| glucosamine | HMDB0001514 | 1.531 | 0.066 | 1.724 | **0.002** |
| oxalic acid | HMDB0002329 | 1.024 | 0.919 | 0.929 | 0.715 |
| threose | HMDB0002649 | 0.772 | 0.414 | 1.043 | 0.772 |
| n-acetyl-tryptophan | HMDB0013713 | 1.076 | 0.797 | 1.220 | 0.354 |
| acetohydroxamic acid | HMDB0014691 | 1.186 | 0.532 | 1.187 | 0.353 |
| aspartyl-glutamate | HMDB0028752 | 0.581 | 0.162 | 0.576 | 0.096 |
| leucine | HMDB00687 | 1.077 | 0.797 | 1.265 | 0.152 |

**Supplemental Table 5.** Pre-screening checklist. A response of “yes” to any of the following resulted in exclusion from the study.

|  |  |  |
| --- | --- | --- |
| Pre-Existing Symptoms Checklist | **INSTRUCTIONS:** Review this list at each visit. If any symptom is present at Baseline, be sure to report it on the ***on medical history*** |  |
| **Symptom** | ☐Yes ☐No | Comments |
| 1. pregnant or breastfeeding | ☐Yes ☐No |  |
| 2. have a bleeding disorder | ☐Yes ☐No |  |
| 3. allergy to the local anesthetic lidocaine | ☐Yes ☐No |  |
| 4. history of stroke, seizures, Parkinson's disease, history of head injury with loss of consciousness, or other dementing disorder | ☐Yes ☐No |  |
| 5. history of alcoholism or drug abuse | ☐Yes ☐No |  |
| 6. History of schizophrenia or currently suffer from bipolar disorder or major depression. | ☐Yes ☐No |  |
| 7. vision or hearing loss severe enough to interfere with cognitive testing | ☐Yes ☐No |  |
| 8. Taking Beta Blockers (ex. Sectral, Tenormin, Zebeta, Lopressor, Corgard, Bystolic, Inderal LA, InnoPran XL) | ☐Yes ☐No |  |
| 9. Taking Neuroleptics (ex. Clozaril, Saphris, Zeprexa, Seroquel) | ☐Yes ☐No |  |
| 10. Taking Narcotic Analgesics (ex. Codeine, Zohydro ER, Oxycodone, Methadone, Hydromorphone, Morphine, Fentanyl) | ☐Yes ☐No |  |
| 11. Taking Anti-Parkinsonian Agents (ex. Sinemet, Symmetrel, Artane, Cogentin, Elderpryl, Azliect, Comtan) | ☐Yes ☐No |  |
| 12. Taking CNS-Active antihypertensive agents (ex. Catapres, Kapvay, Intuniv, Tenex) | ☐Yes ☐No |  |
| **Information obtained by:** |  |  |

**Supplemental Fig. 1.**
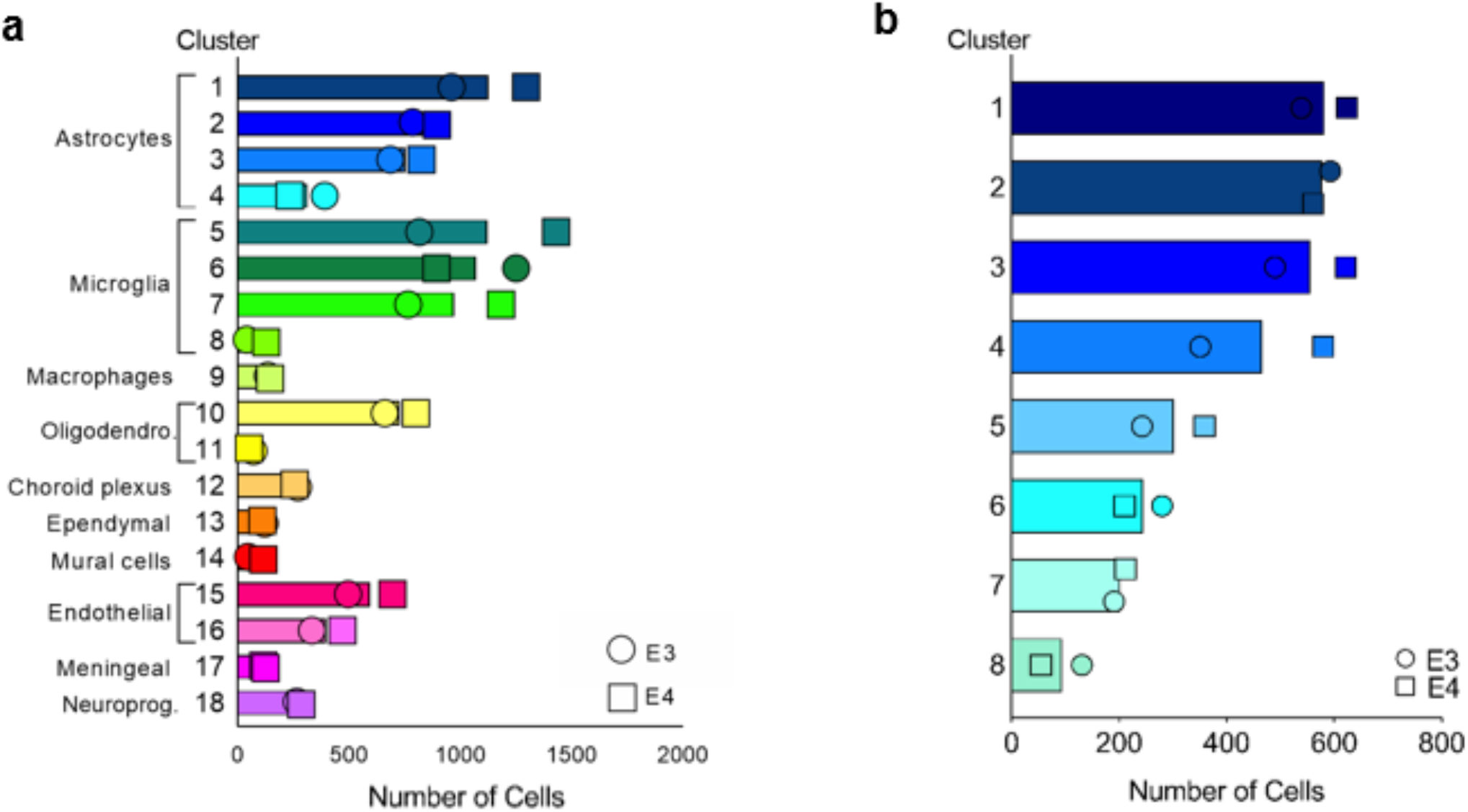
Cluster cell counts. Number of cells in each graph-based cluster from all cells **(a)**, and astrocytes only **(b)**. Bars represent mean number of cells in each cluster, with the number of E3 cells (circles) and E4 cells (squares) noted by symbols.

**Supplemental Fig. 2.**
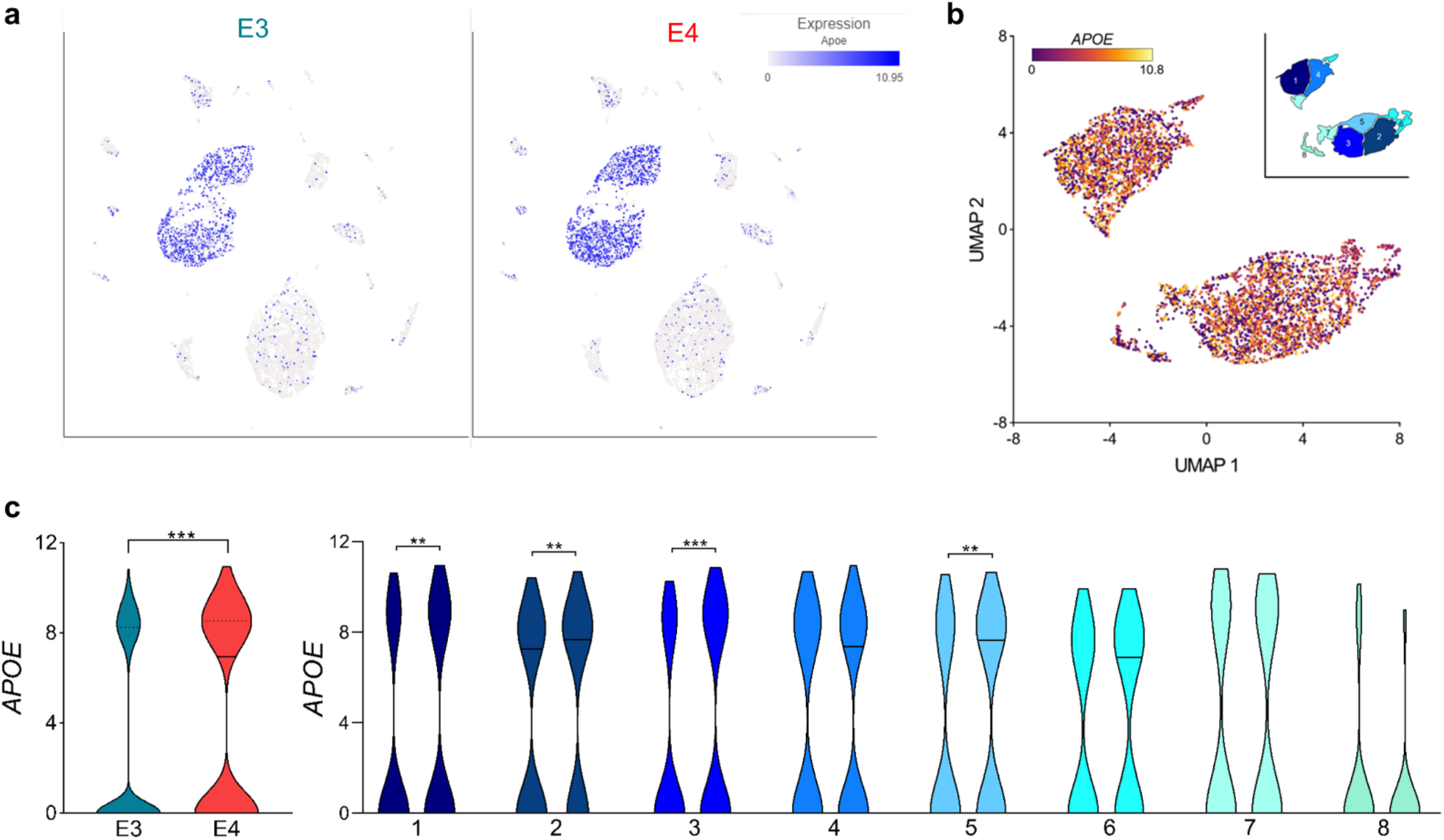
*APOE* expression in single-cells and specific astrocyte clusters. (a) UMAP visualization of E3 (left) and E4 (right) cells showing expression of *APOE. APOE* expression is primarily limited to cells identified as astrocytes. (b) Expression of *APOE* in astrocyte-only UMAP (*Aldoc*+ cells). Inset shows the 8 distinct astrocyte clusters. (c) Violin plots showing expression of *APOE* in all astrocytes (left) and within each astrocyte cluster (right). (\*\**P*<0.01, ***P<0.001, unpaired *t*-test, two-tailed)

**Supplemental Fig. 3.**
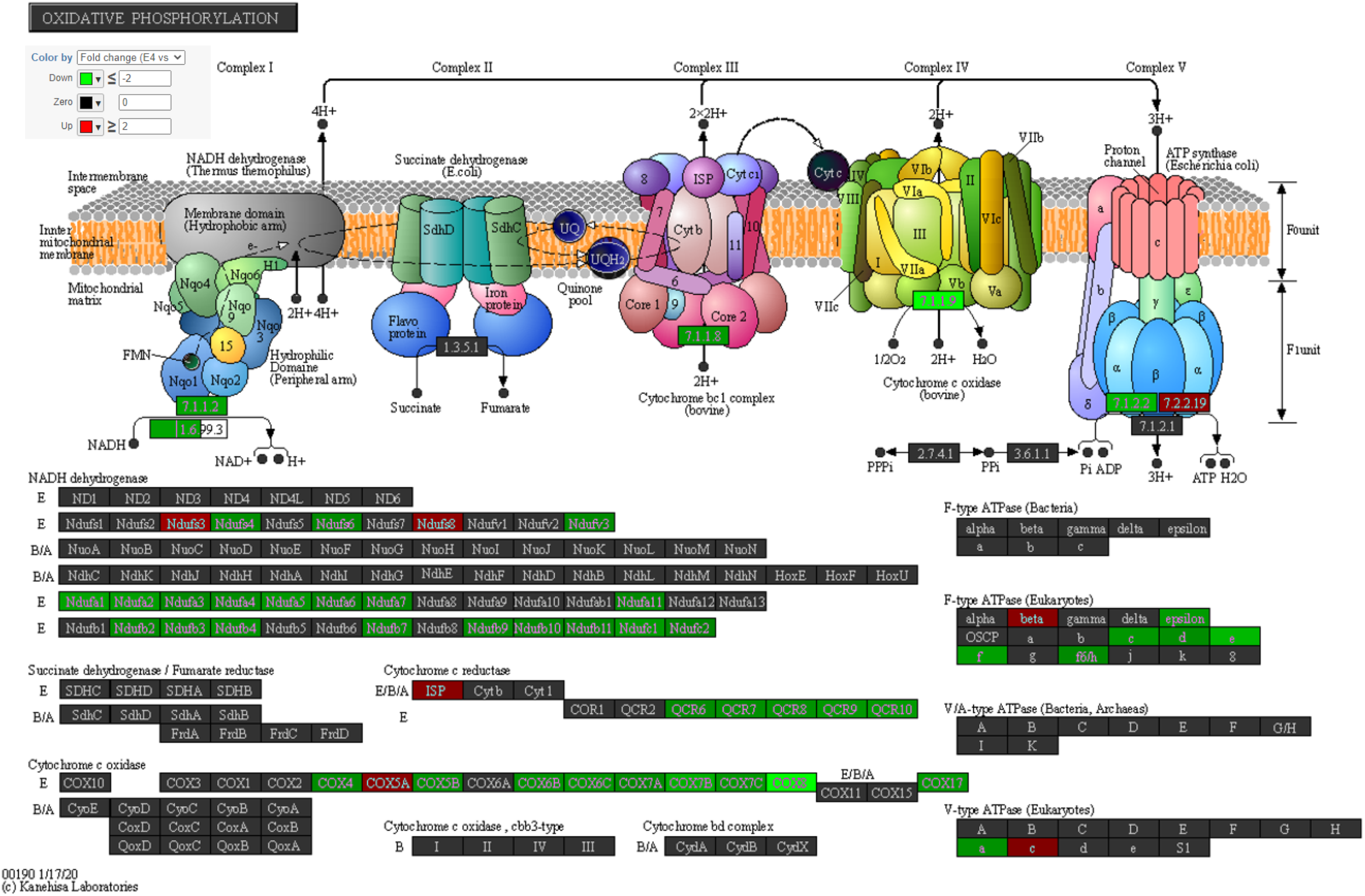
E4 is associated with decreases in many genes of the oxidative phosphorylation KEGG pathway. Pathway map for KEGG pathway “Oxidative Phosphorylation” showing genes differentially expressed between E3 and E4 astrocytes. Genes highlighted in green are downregulated in E4, genes in red are upregulated in E4.

**Supplemental Fig. 4.**
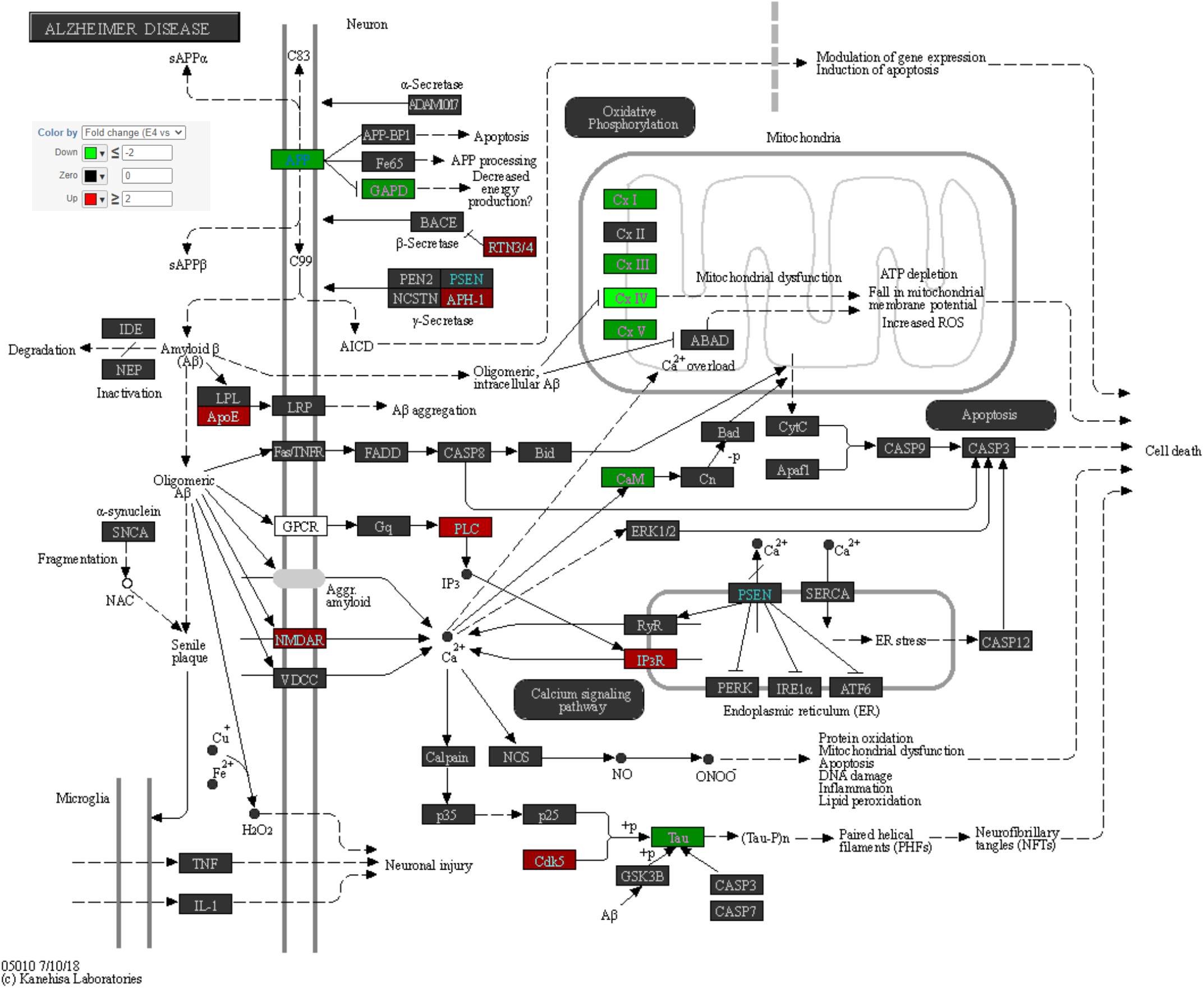
E4 is associated with decreases in many genes of the “Alzheimer’s disease” KEGG pathway. Pathway map for KEGG pathway “Oxidative Phosphorylation” showing genes differentially expressed between E3 and E4 astrocytes. Genes highlighted in green are downregulated in E4, genes in red are upregulated in E4.

**Supplemental Fig. 5.**
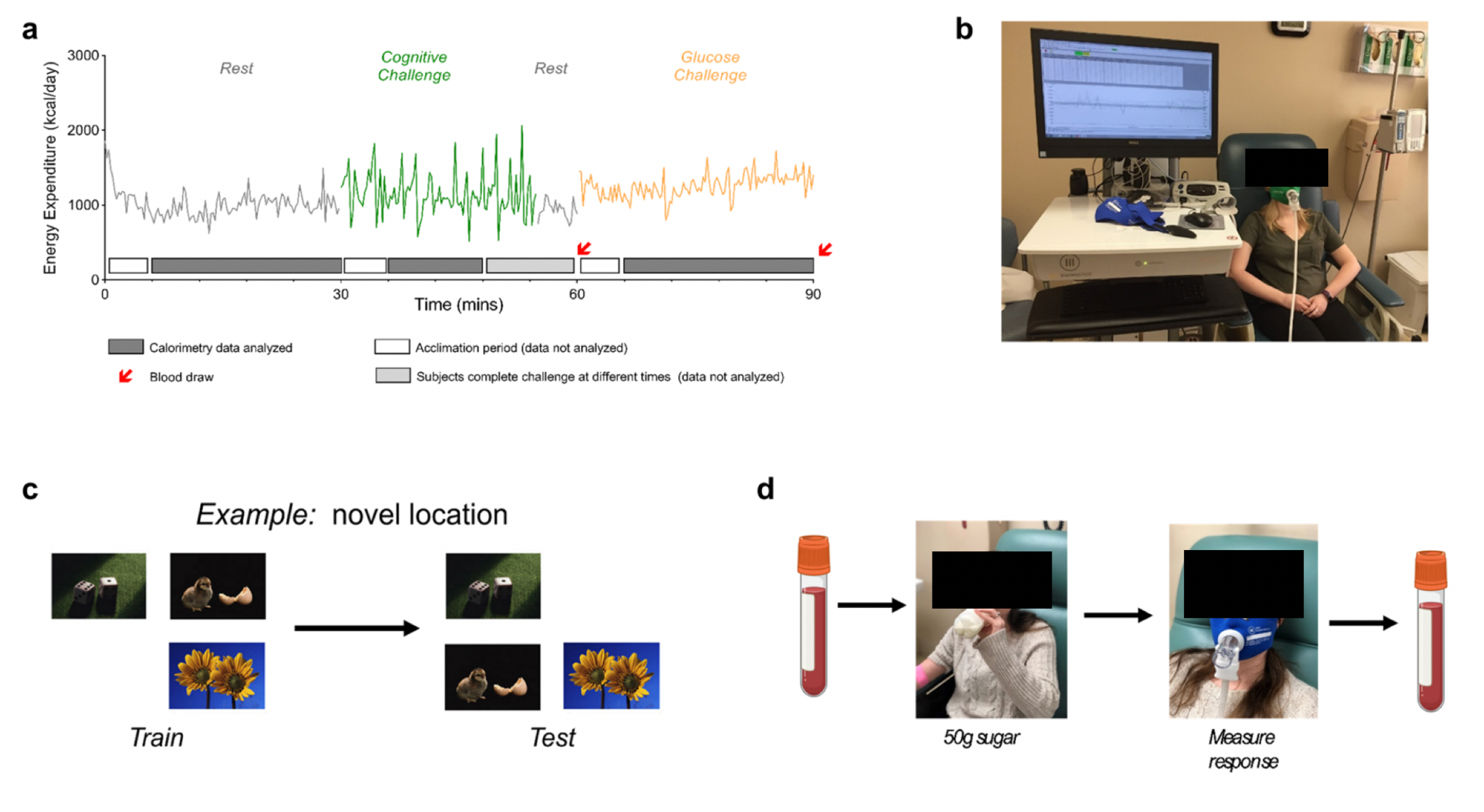
Human indirect calorimetry study design. **(a)** Representative time course of energy expenditure (EE) measures during the three periods of the study (rest in gray, cognitive challenge in green, and glucose challenge in orange). Data was only analyzed during the last 25 minutes of the resting and glucose periods and during a common 5-15 minute span during the cognitive challenge in which all 100 subjects were actively engaged in the task – denoted by grey bar on x axis. Blood was drawn immediately prior and after the glucose challenge. **(b)** Representative photo of a participant during the resting challenge connected to the Ultima MGX indirect calorimetry (IC) system. **(c)** Example slides from the Novel Image Novel Location test used as a cognitive challenge. **(d)** The glucose challenge consisted of a blood draw, followed by ingestion of the 50g sugar drink (all subjects consumed drink within 90 seconds), followed by IC measurement, and a second blood draw.

**Supplemental Fig. 6.**
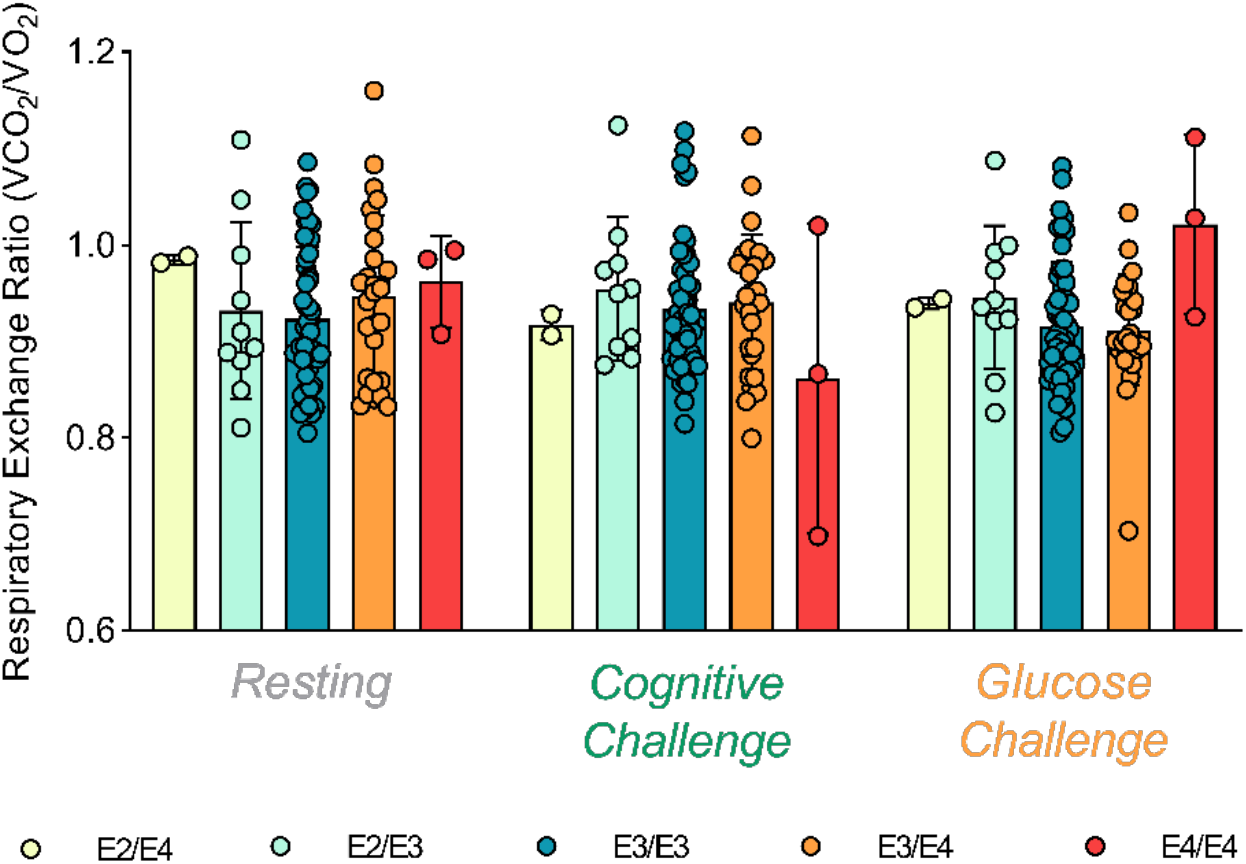
Respiratory Exchange Ratio (RER) does not differ by *APOE* genotype. Respiratory exchange ratio (RER) (VCO_2_/VO_2_) was not significantly different between *APOE* genotypes across any of the three periods tested.

**Supplemental Fig. 7.**
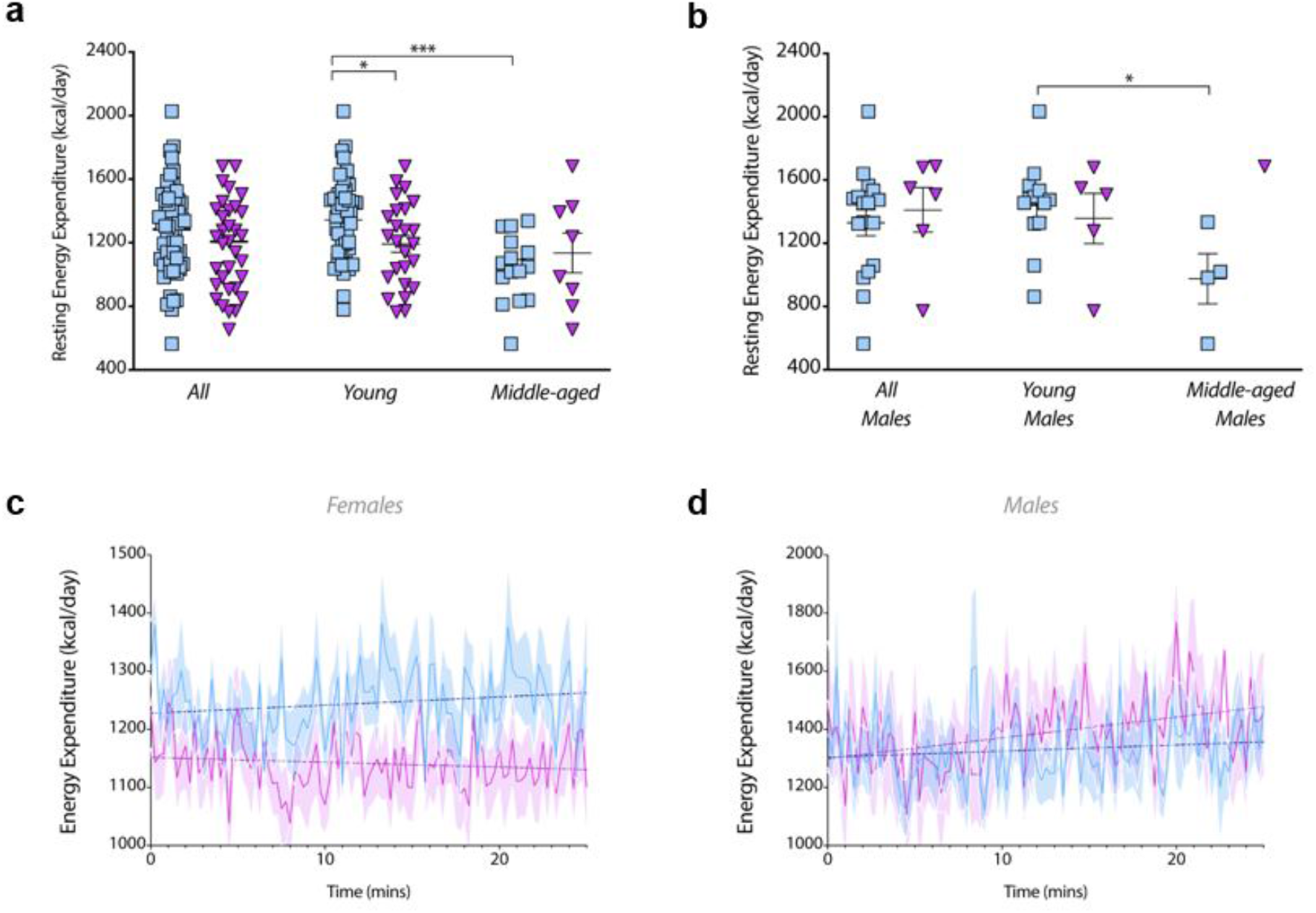
E4 effect on resting energy expenditure. **(a)** E4 non-carriers’ (n=61; blue) and E4 carriers’ (n=33; purple) average resting energy expenditures were determined and stratified by young and middle-aged. (*P<0.05, ***P<0.001, unpaired t-test, two-tailed). **(b)** This was repeated for only male participants (*P<0.05, unpaired t-test, two-tailed; E4-total n=17, young n=13, middle-aged n=4; E4+ total n=6, young n=5, middle-aged n=1). **(c)** Average EE was plotted over the resting period for females and **(d)** males. Dotted lines indicate liner regression results and shaded area are SEMs.

**Supplemental Fig. 8.**
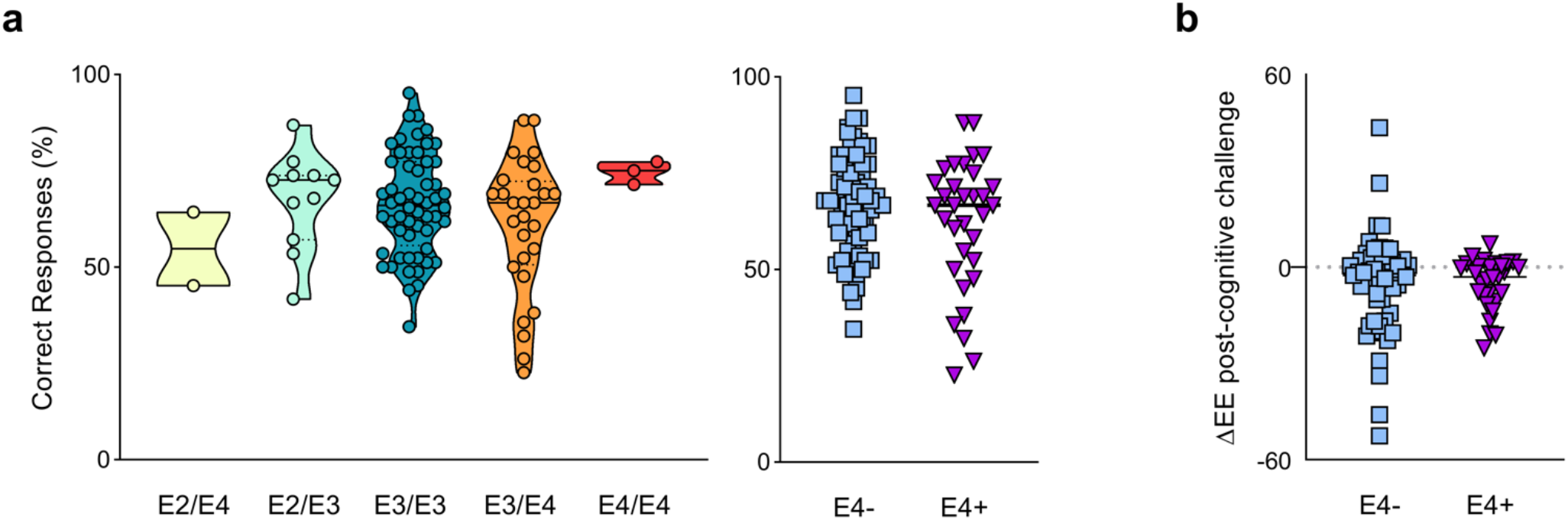
Novel image novel location object recognition test response accuracy by APOE genotype. **(a)** The novel-image-novel-location (NINL) object recognition test contains 7 sets of 12 slides. Each slide has 3 images and 4 possible locations. Each slide is viewed for eight seconds in the order as follows: See Set A, See Set B. Test Set A, See Set C, Test Set B, See Set D, Test Set C, etc. To be considered correct, subjects must identify both the type of change and in which quadrant the change has occurred. The test is designed so that on average subjects answer 60-80% of questions correctly. Total percent correct was calculated for each genotype **(b)** and stratified by E4 carriage. **(c)** Individual slopes of EE after the cognitive challenge showing an average decrease in EE after the challenge.

**Supplemental Fig. 9.**
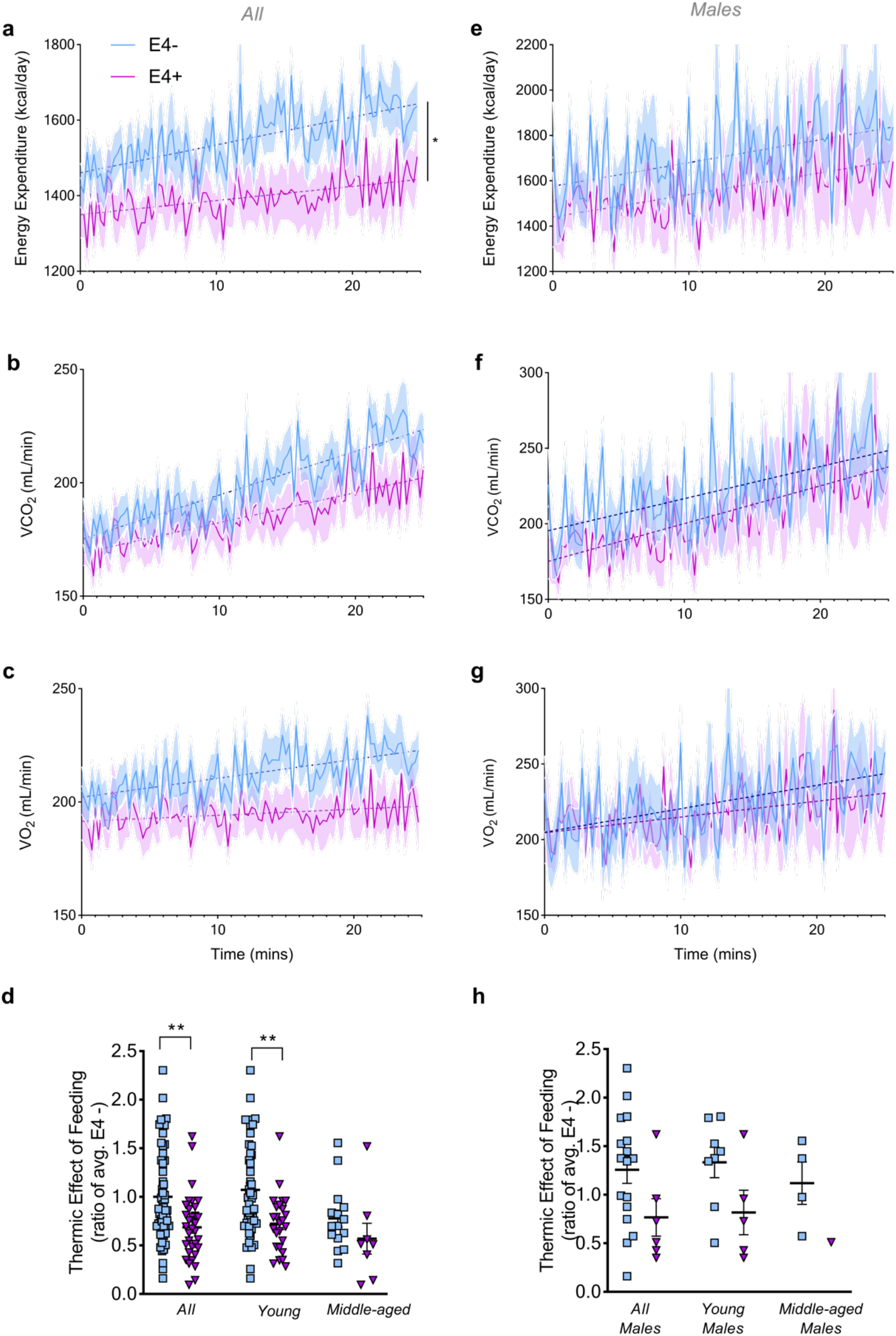
E4 effect on energy expenditure during glucose challenge. **(a)** Energy expenditure **(b)** VCO_2_ and **(c)** VO_2_ was plotted over the glucose challenge period in all E4-(n=61; blue) and E4+ (n=33; purple) participants. (*P<0.05, Two-way ANOVA repeated measures). **(d)** Thermic effect of feeding was determined as a ratio of E4 non-carriers in all, young, and middle-aged participants. (**P<0.01, unpaired t-test, two-tailed) **(e)** Energy expenditure **(f)** VCO_2_ and **(g)** VO_2_ was plotted over the glucose challenge period in male participants (E4-n=17; E4+ n=6). Dotted lines show linear regression trend line, shaded areas refer to SEM. **(h)** Thermic effect of feeding was determined as a ratio of E4 non-carriers in all, young, and middle-aged male participants.

**Supplemental Fig. 10.**
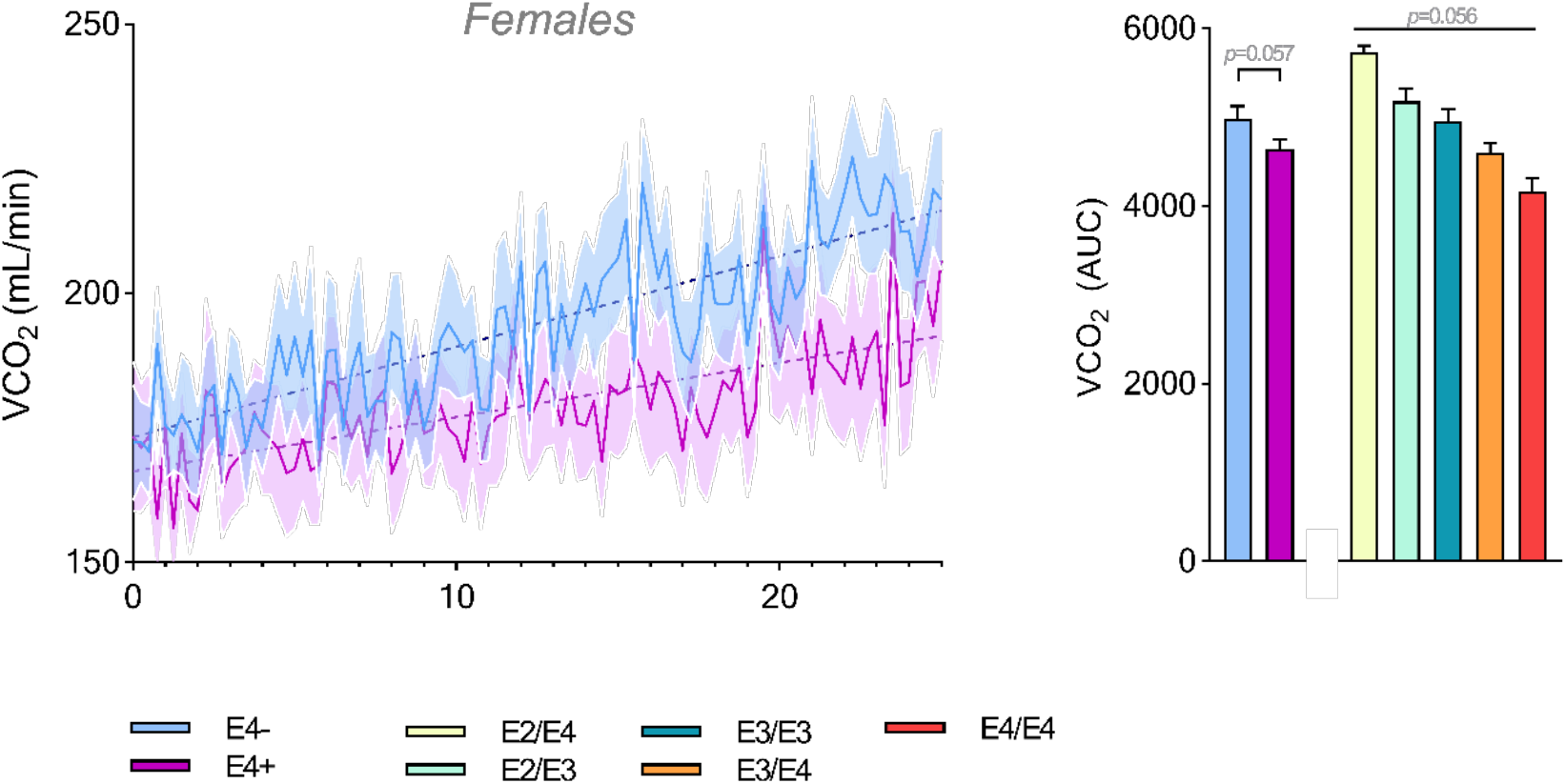
VCO_2_ values during the glucose challenge period. **(a)** Time course of average VCO_2_ values of E4- and E4+ females during the glucose challenge period. Dashed lines refer to linear regression result. **(b)** AUC of VCO_2_ for all participants. (a, Two-way ANOVA repeated measures; b, One-way ANOVA)

**Supplemental Fig. 11.**
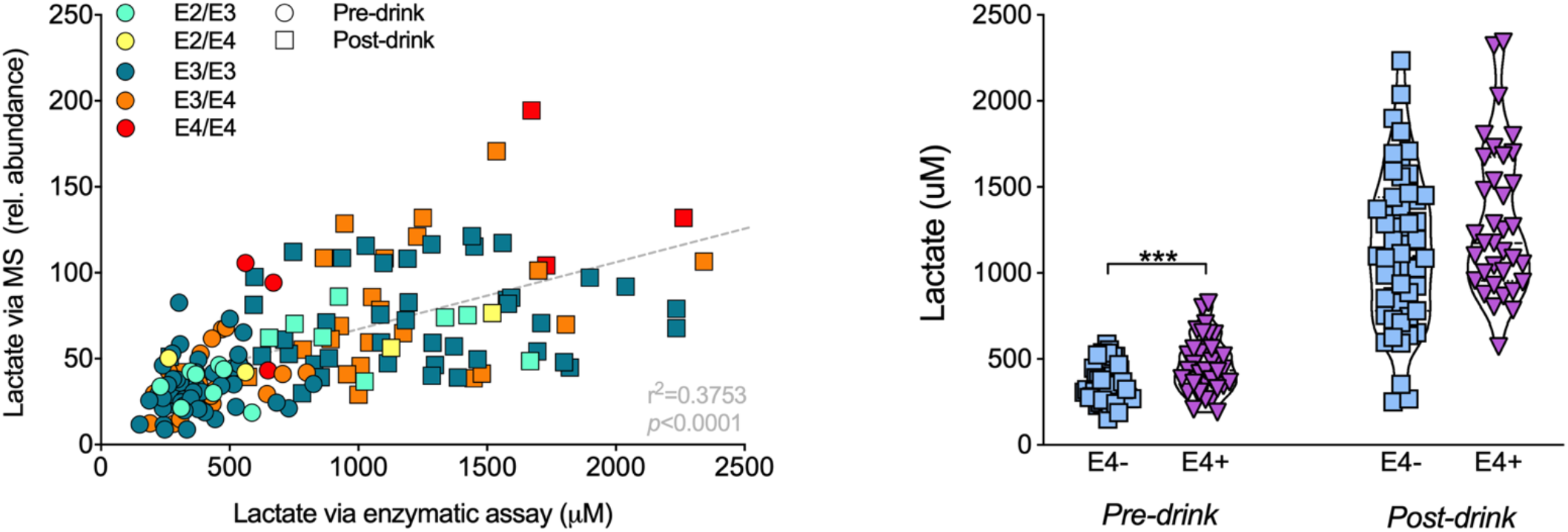
Plasma lactate assessed via enzymatic assay. **(a)** Lactate values quantified by GCMS (relative abundance, y-axis) strongly correlate with lactate values (uM) assessed via enzymatic assay. **(b)** E4 carriers had higher plasma lactate pre-drink and a trend toward higher lactate post-drink (p=0.09) compared to non-carriers, as measured via enzymatic assay.

## Notes

### Competing Interest Statement

The authors have declared no competing interest.

